# Realisation of a key step in the evolution of C_4_ photosynthesis in rice by genome editing

**DOI:** 10.1101/2024.05.21.595093

**Authors:** Jay Jethva, Florian Hahn, Rita Giuliani, Niels Peeters, Asaph Cousins, Steven Kelly

## Abstract

C_4_ photosynthesis is a repeatedly evolved adaptation to photosynthesis that functions to reduce energy loss from photorespiration. The recurrent evolution of this adaptation is achieved through changes in the expression and localisation of several enzymes and transporters that are conventionally used in non-photosynthetic metabolism. These alterations result in the establishment of a biochemical CO_2_ pump that increases the concentration of CO_2_ around rubisco in a cellular environment where rubisco is protected from oxygen thus preventing the occurrence of photorespiration. A key step in the evolution of C_4_ photosynthesis is the change in subcellular localisation of carbonic anhydrase (CA) activity from the mesophyll cell chloroplast to the cytosol, where it catalyzes the first biochemical step of the C_4_ pathway. Here, we achieve this key step in C_4_ evolution in the C_3_ plant *Oryza sativa* (rice) using genome editing. We show that editing the chloroplast transit peptide of the primary CA isoform in the leaf results in relocalisation of leaf CA activity from the chloroplast to the cytosol. Through analysis of fluorescence induction kinetics in these CA relocalisation lines we uncover a role a new role for chloroplast CA in photosynthetic induction. We also reveal that relocalisation of CA activity to the cytosol causes no detectable perturbation to plant growth or leaf-level CO_2_ assimilation. Collectively, this work uncovers a novel role for chloroplast CA in C_3_ plants, and demonstrates that it is possible to achieve a key step in the evolution of C_4_ photosynthesis by genome editing.

**Significance statement:** C_4_ photosynthesis is a highly efficient adaptation to photosynthesis that fuels the world’s most productive crop plants. It evolved from conventional C_3_ photosynthesis through a series of changes in leaf biochemistry and anatomy. Here we achieve a key evolutionary step on the path to C_4_ photosynthesis in rice using genome editing. Specifically, we alter the primary location of carbonic anhydrase activity in the rice leaf from the chloroplast to the cytosol. In doing so, we uncover a novel role for carbonic anhydrase in facilitating the rapid induction kinetics of photosystem II, and initiate a new era of C_4_ engineering using precision breeding techniques.

## Introduction

Photosynthesis is the process by which almost all carbon enters the biosphere. Most terrestrial plants carry out C_3_ photosynthesis, in which CO_2_ is directly fixed into 3-phosphoglycerate (3-PGA) by ribulose-1,5-bis-phosphate carboxylase/oxygenase (rubisco). However, the efficiency of photosynthesis is decreased by the ability of rubisco to catalyse a competing reaction with O_2_. The 2-phosphoglycolate that is produces by this reaction inhibits primary metabolism and must be recycled by photorespiration. The combined opportunity costs and recycling costs attributed to photorespiration result in losses of up to ∼50% of leaf energy under current atmospheric conditions (Walker et al., 2016). Thus, substantial energetic incentives exist for terrestrial plants to evolve adaptations that minimise the occurrence of photorespiration.

To date, over 120 different land plant lineages have evolved biochemical and anatomical adaptations that increase the concentration of CO_2_ compared to O_2_ around rubisco and thus reduce the occurrence of photorespiration (Gilman et al., 2023; Sage, 2016). These evolutionarily distinct solutions to the photorespiration problem can be categorised into two different photosynthetic types: Crassulacean Acid Metabolism (CAM) and C_4_ photosynthesis. In both altered versions of photosynthesis, plants do not directly fix CO_2_ into 3-PGA, instead they utilise carbonic anhydrase (CA) to produce bicarbonate from CO_2_ and then use a series of biochemical reactions and transmembrane transport steps to deliver and concentrate this CO_2_ in a cellular environment where rubisco is compartmentalized. This concentrated CO_2_ is then fixed into 3-PGA by rubisco with limited reactions with oxygen causing a substantial decrease in production of 2-phosphoglycolate. Consequently, plants that operate biochemical CO_2_ concentrating mechanisms have reduced photorespiratory energy loss, and corresponding increases in radiation, nitrogen and water use efficiencies (Brown, 1978; Way et al., 2014; Zhu et al., 2008)

The advantages of operating a CO_2_ concentrating mechanism, coupled with the apparent reproducibility with which nature has implemented these mechanisms in disparate plant lineages, has motivated international efforts to engineer CO_2_ concentrating mechanisms into C_3_ plants (Hibberd et al., 2008; von Caemmerer et al., 2012). Although there are differences in the implementation of CO_2_ concentrating mechanisms between species, they all share one common evolutionary innovation - the relocalisation of CA activity from the chloroplast in C_3_ plants (Badger & Price, 1994) to the cytosol in both CAM and C_4_ plants (Badger, 2003; Hatch & Burnell, 1990). To date, the evolutionary mechanisms through which this relocalisation occurs has only been investigated in two C_4_ species, *Flaveria bidentis* (Tanz et al., 2009) and *Neurachne munroi* (Clayton et al., 2017). In both cases, the relocalisation of CA activity from the chloroplast to the cytosol occurred through mutation in the chloroplast transit peptide of the isoform of CA that is highly expressed in the chloroplast, rather than by upregulation of a CA isoform that is localised to the cytosol. Thus, while other ways to achieve a shift in location in CA activity may exist, there is precedent from two independent origins of C_4_ photosynthesis in both eudicots and monocots that relocalisation of CA activity to the cytosol is achieved through alteration of organellar targeting of the already highly expressed chloroplast localised isoform of CA.

In C_3_ plants, CA exists in three subtypes (α-CA, β-CA, γ-CA) with the chloroplast localised β-CA being the most abundant in leaves (Ludwig, 2016; Okabe et al., 1984; Poincelot, 1972; Tsuzuki et al., 1985). Although a diverse set of functions have been proposed for different CA’s in C_3_ photosynthesis, it is generally thought that the primary role is to help maintain CO_2_ levels for rubisco carboxylation (Badger & Price, 1994). However, several studies have revealed limited impact on photosynthesis when CA activity is reduced (Ferreira et al., 2008; Majeau et al., 1994; Price et al., 1994), perhaps because of its high turnover rate of *k*_catCO2_ ≈ 10^6^ s^−1^ (Lindskog & Coleman, 1973). CA may also influence the redox state of photosystem II through the provision of bicarbonate in the chloroplast, as studies suggest bicarbonate binding to PSII’s non-heme iron affects electron transport (Brinkert et al., 2016; Cox et al., 2009). Given that the relocalisation of CA activity from the chloroplast to the cytosol is a recurrent feature of both C_4_ and CAM photosynthesis, the question arises as to what are the consequences of this relocalisation on function of photosystem II, and on CO_2_ assimilation and plant growth during C_4_ evolution.

Here we followed precedent from evolution to achieve the relocalisation of CA activity from the chloroplast to the cytosol in the C_3_ plant *Oryza sativa* (rice) using genome editing. Specifically, we targeted the chloroplast transit peptide of chloroplast localised CA to create both deletion and frame shift mutations to relocalise CA activity from the chloroplast to the cytosol. We demonstrate that genome-edited lines with relocalised CA activity exhibit no deleterious effects on leaf-level CO_2_ assimilation or growth and uncover a novel role for CA in the function in maintaining high rates of electron transfer in photosystem II. Collectively, these results provide new insight in the function chloroplast CA in C_3_ plants, and shown that it is possible to recreate a key step in the evolution angiosperm carbon concentrating mechanisms using genome editing.

## Results

### Generation of genome edited rice lines with altered chloroplast transit peptides for the primary carbonic anhydrase

To determine the relative abundance of each carbonic anhydrase (CA) isoform in mature rice leaves, and to confirm the gene model of the chloroplast localised isoform for genome editing, a dataset of mature leaf RNA-Seq (Ermakova et al., 2021) was evaluated in the context of the experimentally defined transcription start sites (Murray et al., 2022). Evaluation of these data identified the gene OsKitaake01g256600 as the most abundant expressed CA in the leaf, accounting for more than 98% of all transcripts encoding CA enzymes (Supplemental File 1, Figure S1). Manual evaluation and revision of the genome annotation for this gene in the context of the RNA-Seq data identified 3 distinct transcripts produced from two discrete transcription start sites in mature rice leaves (Figure 1A). The most abundant transcript from this gene contained a chloroplast transit peptide, while the other two transcripts from the locus lacked targeting sequences and were predicted to encode CA proteins that are localised to the cytosol (Figure 1B, Supplemental File 1 Figure S2 and Figure S3)).

**Figure 1.**
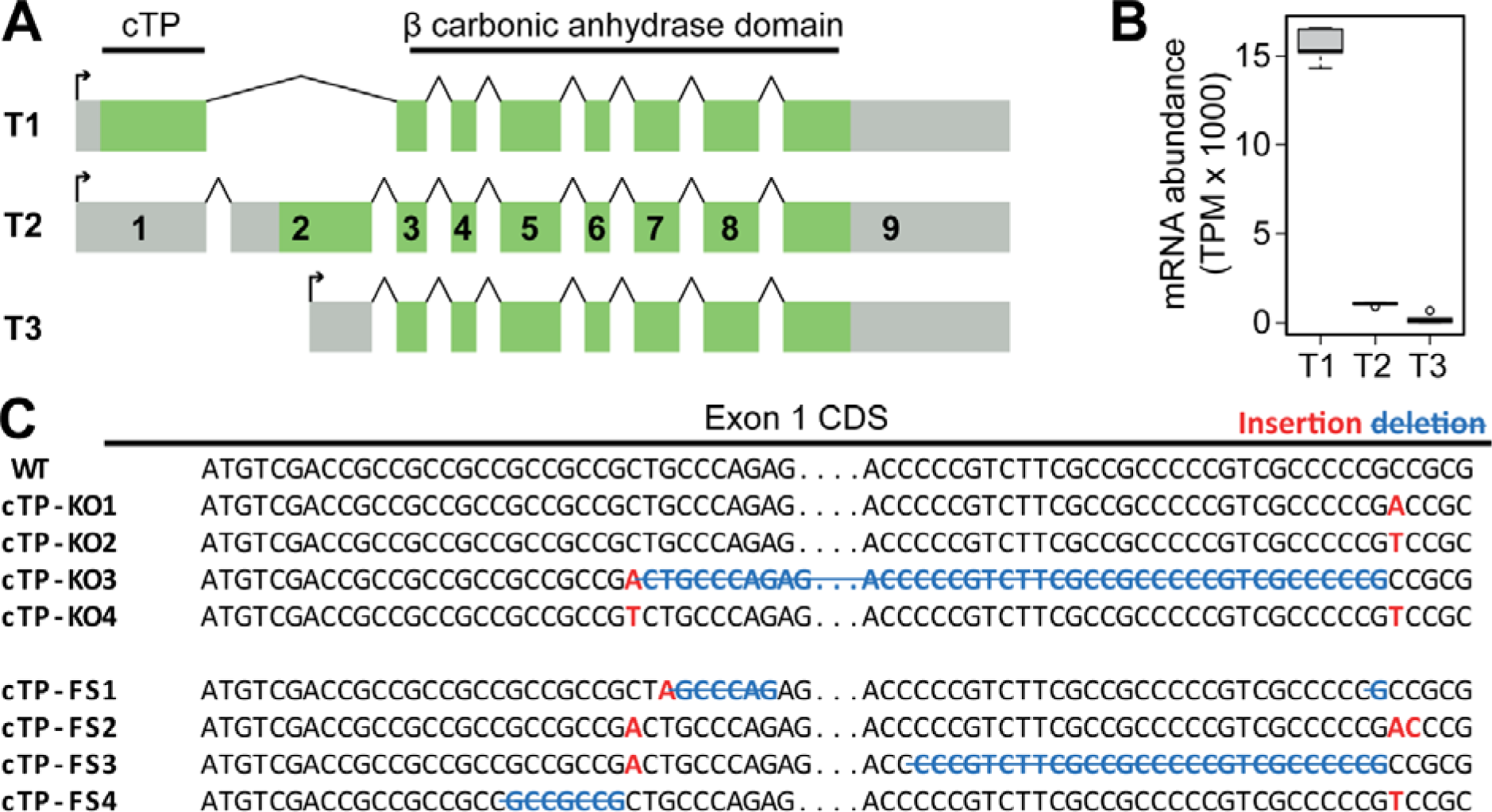
Gene models, mRNA abundance, and genome edited alleles for the primary leaf carbonic anhydrase OsKitaake01g256600. A) Gene model diagram depicting the three transcript variants produced from the OsKitaake01g256600 locus through a combination of alternative splicing and alternative transcription start site use. Transcription start sites depicted by an arrow. Exons (green boxes) and UTRs (grey boxes) are to scale, introns (connecting lines) are not to scale. Exon numbers indicated on transcript 2. B) mRNA abundance of the three transcript variants in mature rice leaves. C) The 8 different CRISPR alleles generated and evaluated in this study.

To attempt to mimic evolution and relocalise CA activity from the chloroplast to the cytosol, a set of guide RNAs were designed to target the chloroplast transit peptide encoded in exon 1 (Supplemental File 1, Figure S2). Guides were designed at the start and the end of coding sequence for the transit peptide to attempt generate a series of mutants with altered transit peptide sequences. Several rounds of transformation and screening by sequencing resulted in the identification of eight independent genome edited lines that were predicted to lack chloroplast localised CA activity (Figure 1C). Four of these lines had mutations that caused the introduction of premature termination codons before the CA domain in transcript 1, but did not affect the sequences of transcript 2 or 3. These lines are collectively referred to as the “chloroplast transit peptide knock out” (cTP-KO) lines (Supplemental File S2, Figure 1C) as carbonic anhydrase activity should be lost from the chloroplast but retained in the cytosol. In addition, four lines were generated in which the chloroplast transit peptide was knocked out of frame but the entire CA domain remained in frame and unaffected. This was achieved by the insertion or deletion of one or more bases at the start of the chloroplast transit peptide, followed by a restoration of the reading frame at the end of the chloroplast transit peptide by a complimentary mutation (Supplemental File S2, Figure 1C). These lines are collectively referred to as the “chloroplast transit peptide frame shift” (cTP-FS) lines.

### CA isoforms with frame-shifted transit peptides localise to the cytosol in rice mesophyll protoplasts

To determine whether the novel cTP-FS coding sequences created by editing the chloroplast transit peptide resulted in a change in subcellular localisation of the encoded protein, the edited sequences were subject to computational subcellular localisation prediction. While the proteins encoded by both the cTP-FS2 and cTP-FS3 alleles were predicted to encode cytosolic enzymes (Supplemental File 1, Figure S2), the proteins encoded by cTP-FS1 and cTP-FS4 were predicted to have weak targeting to the chloroplast (Supplemental File 1, Figure S3). Thus, to confirm or refute these computational predictions the coding sequences of all cTP-FS alleles were fused to mTurquoise2 and transiently expressed in rice mesophyll protoplasts (Figure 2A). Analysis of the localisation of the fusion proteins revealed, that in contrast to the computational prediction, each cTP-FS allele localised the cytosol with no detectable localisation to the chloroplast (Figure 2B). Each fusion protein produced the same localisation pattern as free monomeric mTurquoise2 (Figure 2B) and cytosolic CA (with no transit peptide) fused to mTurquoise2 (Figure 2B). The coding sequence contained in transcript variant 1, that contains the non-edited wild-type transit peptide sequence, was also evaluated to confirm the chloroplast localisation of the primary transcript variant (Figure 2B). Thus, despite weak computational predicted chloroplast targeting for cTP-FS1 and cTP-FS4, all four cTP-FS alleles localise to the cytosol in rice mesophyll cells. Accordingly, CA activity arising from all four cTP-FS edited alleles should now be localised in the cytosol. As all cTP-FS lines and all cTP-KO lines appeared equivalent, two independent lines of each type (cTP-FS1, cTP-FS4, cTP-KO1 and cTP-KO4) were taken forward for subsequent analyses.

**Figure 2.**
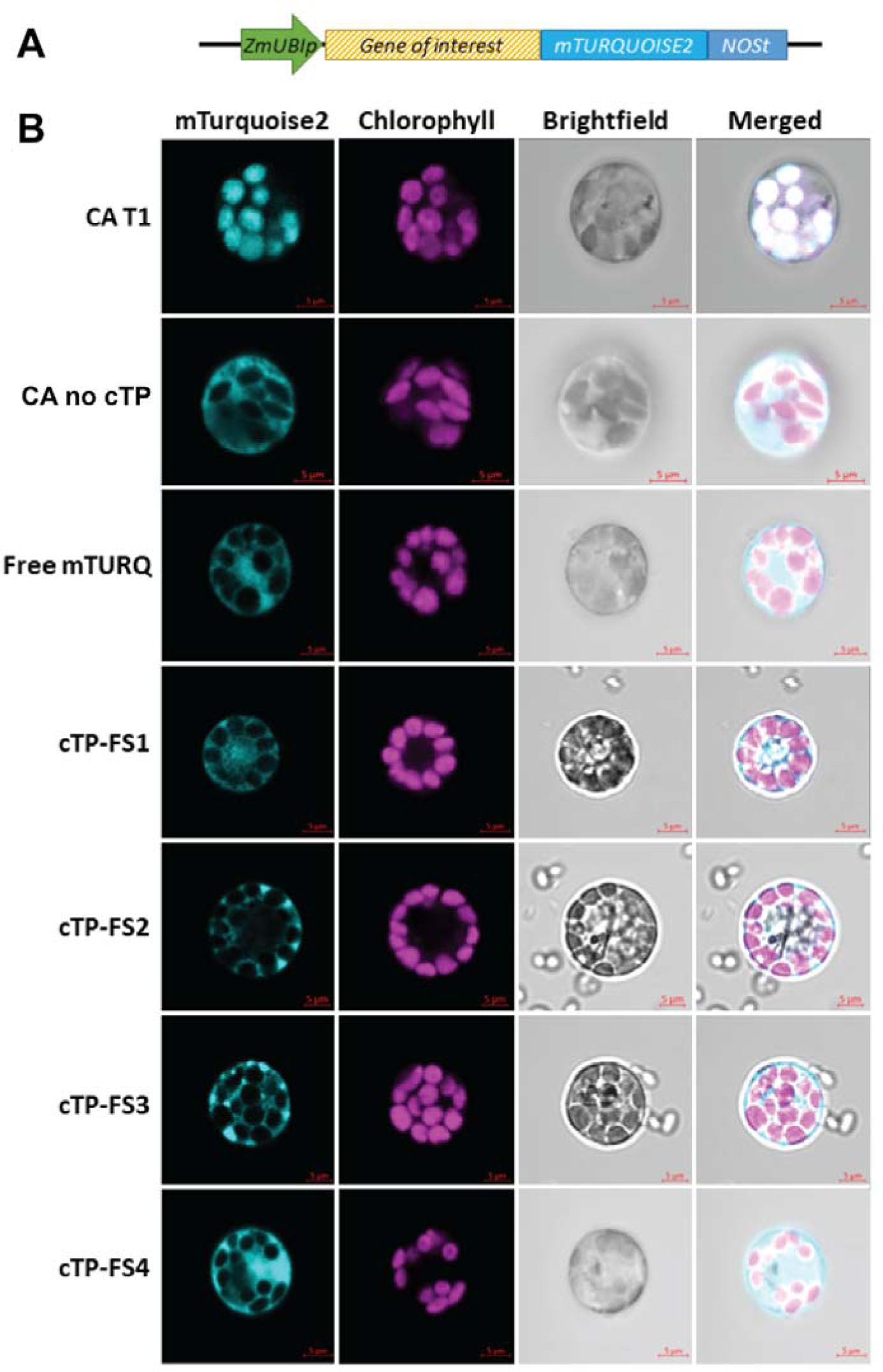
Subcellular localisation of cTP-FS carbonic anhydrase alleles fused to mTurquoise2 in rice protoplasts. A) Schematic representation of expression construct. ZmUBIp is the *Zea mays* ubiquitin promoter. NOSt is the nopaline synthase terminator. B) Confocal microscope images of mTurquoise2 fusion protein fluorescence. The four cTP-FS alleles are shown alongside the wild-type transcript 1 (CA T1), wild-type transcript 2 (CA T3) and mTurquoise2 not fused to any coding sequence (Free mTURQ). Chlorophyll autofluorescence is shown to indicate the location of the chloroplasts.

### Genome edited lines have no detectable CA activity in the chloroplast with all remaining CA activity localised in the cytosol

Given that the cTP-KO alleles no longer encoded a chloroplast localised CA, and that all of the cTP-FS alleles had relocalised the chloroplast CA to the cytosol, it was next investigated whether these sequence changes resulted in a change in the localisation of CA activity within the cell. To evaluate this, the genome edited lines were first subject to transcriptome sequencing to understand how the introduced edits altered the mRNA abundance for the three transcripts originating from the two distinct promoters in the locus. While the mRNA abundance of transcript 2 and transcript 3 (both encoding cytosolic isoforms of CA) were unaffected by the genome edits (Figure 3A, Supplemental File 3), transcript abundance from transcript 1 (encoding the chloroplast targeted isoform of the gene) were significantly downregulated compared to wild-type levels (Figure 3A, p < 10^−10^ DESeq2, Supplemental File 3). This reduction was more pronounced in the cTP-KO (92% reduction in mRNA abundance) than in the cTP-FS lines (77% reduction in mRNA abundance), presumably due to the premature termination codon in the first exon of the transcript leading to nonsense mediated decay of the transcript (Figure 3A, p < 10^−10^ DESeq2, Supplemental File 3). Importantly, there was also no effect on the mRNA abundance of any other gene encoding a CA (Supplemental File 4). Thus, introduction of small nucleotide changes by genome editing, irrespective of whether they introduced a mis-sense or non-sense mutation, resulted in dramatic reduction in the abundance of transcript 1 with no effect on the abundance of transcript 2 or 3.

**Figure 3.**
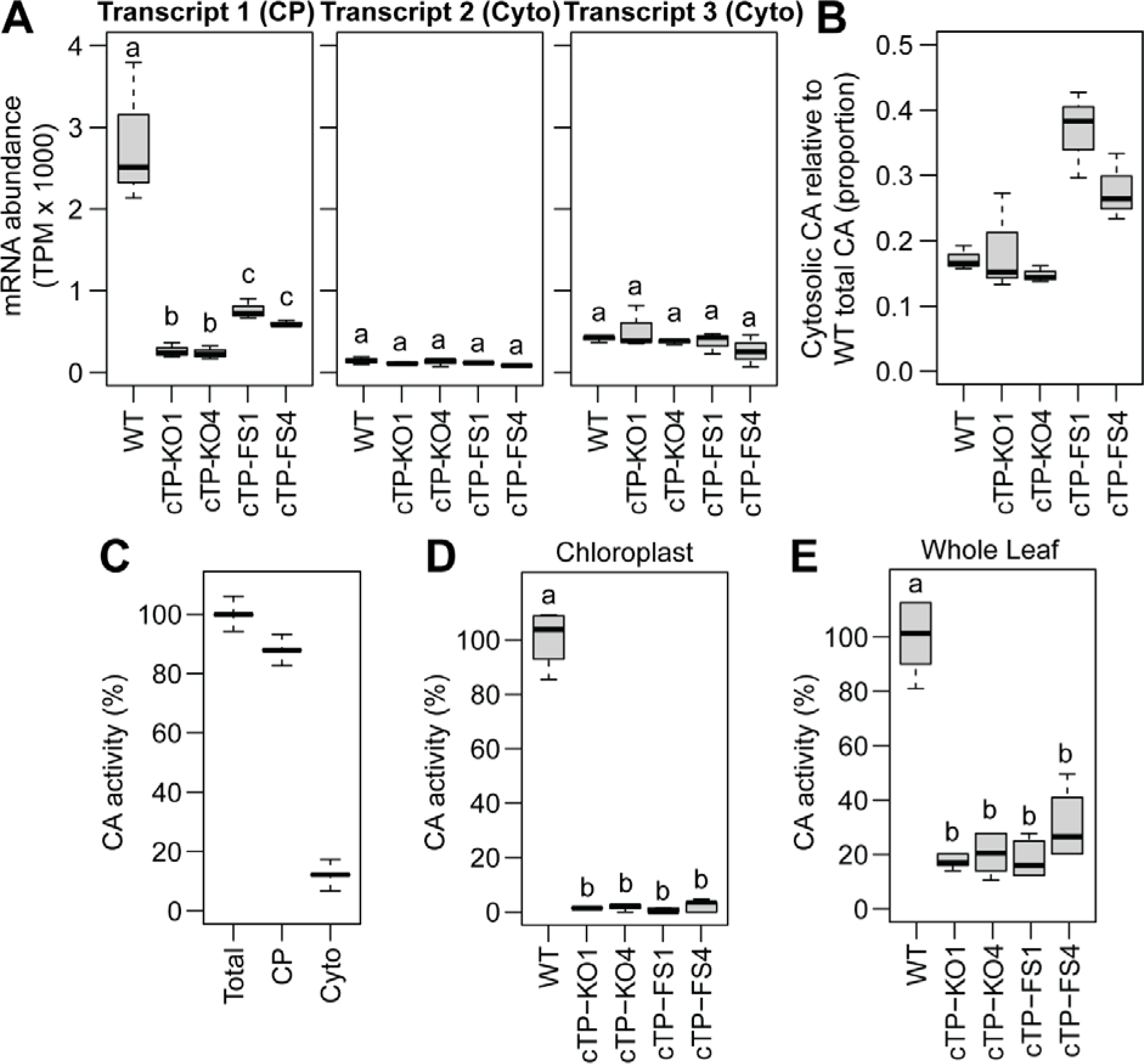
CA transcript abundance and CA activity of WT and genome edited lines. A) The mRNA abundance of the three transcript variants in each of the genome edited and wild-type lines. CP: chloroplast targeted. Cyto: Cytosol targeted. B) The abundance of cytosolic encoded transcripts compared to WT total CA transcript abundance. C) The CA activity contained within the whole leaf and chloroplast fraction of wild-type plants. D) The CA activity contained with the chloroplast of wild-type and genome edited plants, normalised to chloroplast activity. E) The CA activity contained within the whole leaf of wild-type and genome edited plants, normalised to wild-type whole leaf activity. Letters above boxplots represent statistically significant differences (*p* ≤ 0.01) in mean values as determined by Fisher LSD post-hoc analysis following a one-way ANOVA.

The transcriptome data above indicate that CA activity should now be absent from the chloroplast and exclusively localised in the cytosol. To estimate the relative level of CA activity that is present in the cytosol of the genome edited lines, the abundance of transcripts encoding cytosolic CA isoforms was summed and compared to the total transcript abundance for CA normally present in wild-type rice leaves. This revealed that transcripts encoding cytosolic CA accumulated to an equivalent of 15-37% of total wild-type CA transcript abundance in the genome edited lines (Figure 3B). Equivalent to or larger than the proportion of transcripts in wild-type plants that encode cytosolic CA (17%, Figure 3B). Thus, while transcripts encoding chloroplast localised CA are absent, transcripts encoding cytosolic CA accumulate to substantial levels in the genome edited lines.

To demonstrate that the patterns of transcript accumulation observed in the genome edited plants resulted in a corresponding change in localisation of enzyme activity, the genome edited plants were subject to CA enzyme activity assays. In wild-type plants 17% of CA activity is present in the cytosol while 83% is found in the chloroplast (Figure 3C). However, when chloroplasts were isolated from leaves of the cTP-FS or cTP-KO lines there was no detectable CA activity in the chloroplasts (Figure 3D). However, consistent with the transcriptome the genome edited lines exhibited 17-30% of wild-type CA activity (Figure 3E). Given that all expressed CA isoforms are predicted to encode cytosol localised CA, that there was no upregulation of any other CA enzyme, and that there was no detectable activity in the chloroplast, the genome editing had successfully removed CA activity from the chloroplast and all CA activity now resides in the cytosol. Henceforth, the genome edited lines are collectively referred to as the CA relocalisation lines.

### Relocalisation of CA activity form the chloroplast to the cytosol does not result in differential expression of C_4_ cycle genes or substantial changes to the transcriptome

Given that we had mimicked a key step in the evolution of C_4_ photosynthesis, we sought to investigate whether this change resulted in alteration in expression of any other C_4_ cycle genes. Analysis of the transcriptome data revealed that none of the genes encoding enzymes (Furbank, 2016) or transporters (Mattinson & Kelly, 2024) thought to function in the C_4_ cycle of CAM photosynthesis were differentially expressed as a result of relocalisation of CA activity to the cytosol (Figure 4). Furthermore, transcriptome-wide differential expression analysis identified just 23 genes that were downregulated and 50 genes that were upregulated in the CA relocalisation lines when compared to wild-type plants (Supplemental File 3). There were no significantly overrepresented GO terms in either gene set and manual inspection of the gene list did not identify any differentially expressed genes that function in CO_2_ assimilation or related pathways. Thus, the relocalisation of CA activity from the chloroplast to the cytosol did not induce gene expression changes associated with C_4_ evolution, and resulted in minimal perturbation to the whole leaf transcriptome.

**Figure 4.**
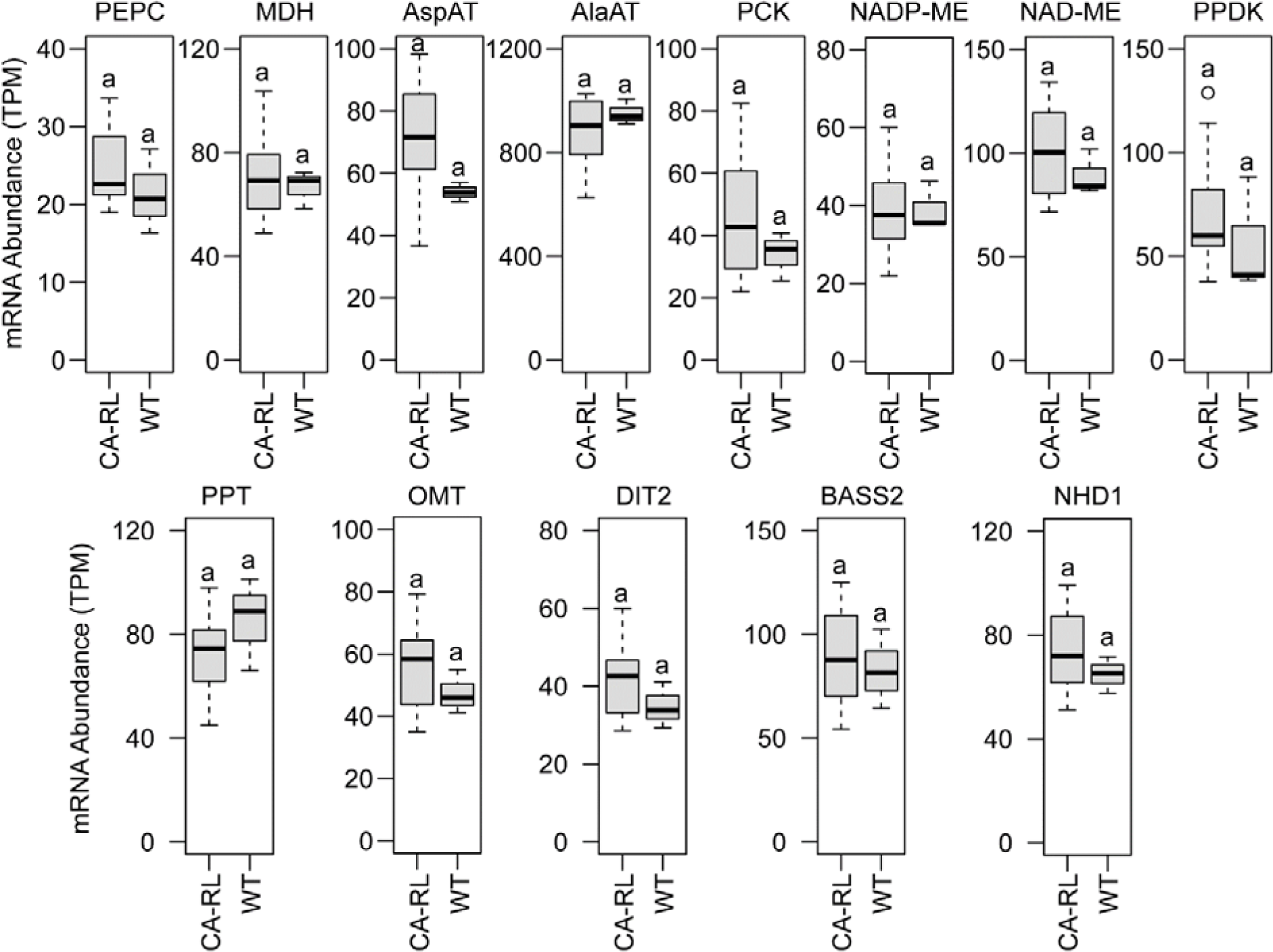
Changes in mRNA abundance of genes encoding enzymes and transporters of the C_4_ cycle. The cTP-FS and cTP-KO lines are collectively shown as CA-RL (CA relocalisation). The name of the enzyme or transporter under consideration is shown above each boxplot. PEPC: Phospho*enol*pyruvate carboxylase. MDH: Malate dehydrogenase. AspAt: Aspartate amino transferase. AlaAT: Alanine amino transferase. PCK: Phospho*enol*pyruvate carboxy kinase. NADP: NADP-malic enzyme. NAD-ME: NAD-malic enzyme. PPDK: Pyruvate, phosphate dikinase. PPT: Phospho*enol*pyruvate/phosphate translocator. OMT: Oxaloacetate/malate transporter. DIT2: Dicarboxylate transporter: BASS2: Bile acide sodium symporter. NHD1: Sodium:hydrogen antiporter 1. Letters above boxplots represent statistically significant differences (*p* ≤ 0.01) in mean values as determined by DESeq 2.

### Growth and CO_2_ assimilation are unaffected in the CA relocalisation lines

Given that we had mimicked a key step in the evolution of C_4_ photosynthesis, we sought to determine whether this change resulted in a deleterious effect on growth or CO_2_ assimilation. Comparison of the CA relocalisation lines and wild-type plants revealed no difference in morphology (Figure 5A), no difference in growth rate (Figure 5B), no difference in final plant height (Figure 5C), and no difference in 100 seed weight (Figure 5D). Consistent with lack of growth defect, there was also no difference in photosynthetic rate (Figure 5E), stomatal conductance to CO_2_ diffusion (Figure 5F) or substomatal CO_2_ concentration (Figure 5G) at either ambient or low O_2_ concentration between the CA relocalisation lines and wild-type plants. Together, these results indicate that under these conditions CO_2_ supply to rubisco has been unaltered. Thus, change in location of CA activity from the chloroplast to the cytosol in rice does not affect plant growth or CO_2_ assimilation.

**Figure 5.**
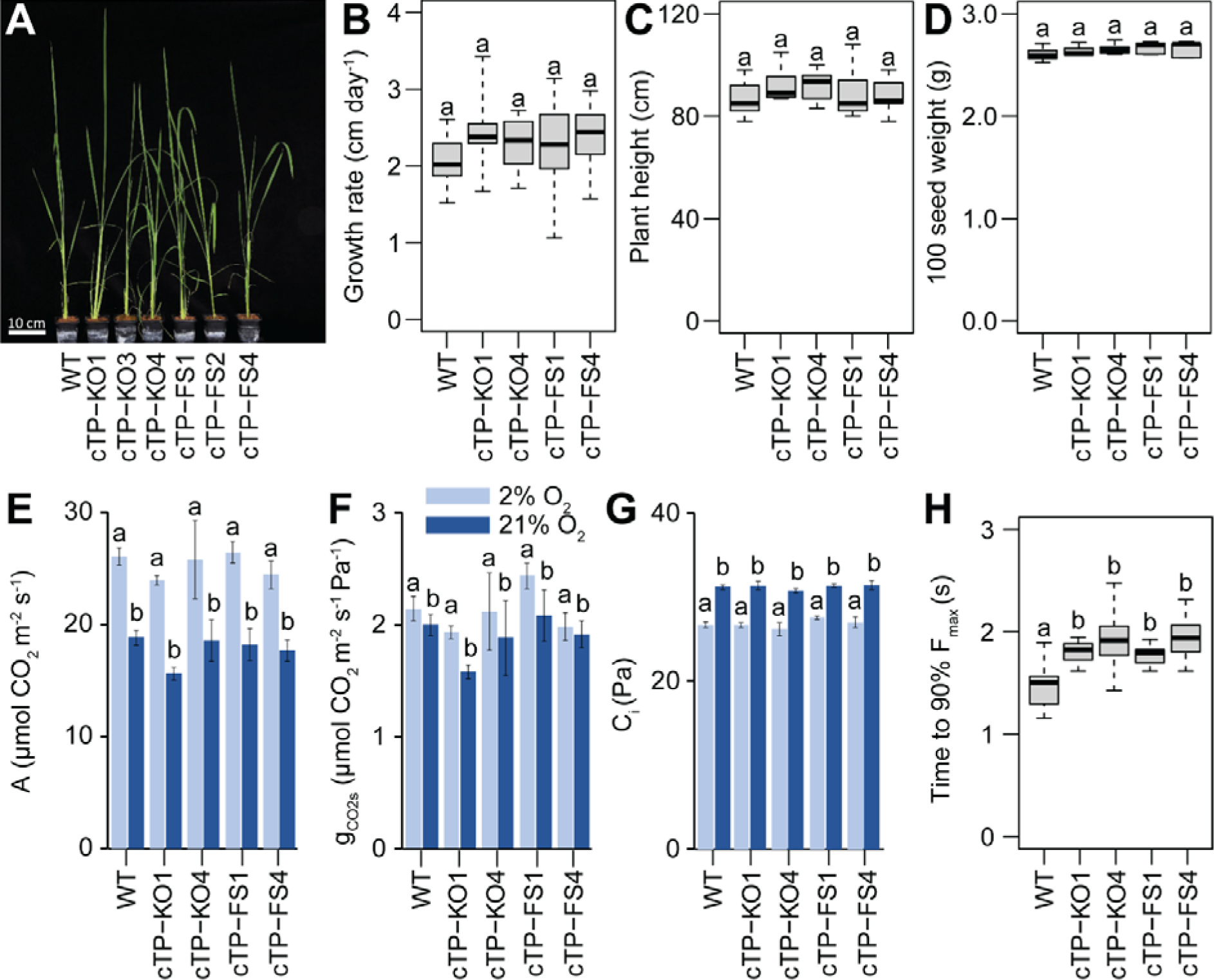
Phenotypic analysis of the CA relocalisation lines. A) Photograph of representative plants at day 25 post transplanting to soil. B) Growth rate calculated from 17-day period from day 8 to day 25 post transplanting to soil. C) Final plant hight measurements taken on day 58 after transplanting to soil. D) 100 seed weight. E) Leaf net photosynthetic rate at 2% and 21% oxygen. F) Stomatal conductance to CO_2_ diffusion at 2% and 21% oxygen. G) CO_2_ partial pressure in the intercellular air space at 2% and 21% oxygen. H) Time to 90% maximal fluorescence. For panels E, F and G Letters above bar plots represent statistically significant differences in mean values as determined by Fisher LSD post-hoc analysis following a two-way ANOVA (factorial design with five plant types and two O_2_ levels; p< 0.05)). For all other plots, letters above boxplots represent statistically significant differences (*p* ≤ 0.01) in mean values as determined by Fisher LSD post-hoc analysis following a one-way ANOVA.

Although there was no perturbation to growth or CO_2_ assimilation, we hypothesised that a lack of CA activity, and by consequence a reduced concentration of bicarbonate in the chloroplast, could influence the function of photosystem II. The rationale for this hypothesis is that bicarbonate plays a role in electron transfer within photosystem II by binding to the non-heme iron positioned at Q_A._ In the presence of bicarbonate the midpoint potential of Q_A_/Q_A_^−^ is −145 mV, while in the absence of bicarbonate it is −70 mV (Brinkert et al., 2016). We posited, that if the CA relocalisation lines had a reduced abundance of bicarbonate in the chloroplast, then this would cause a change in the midpoint potential of Q_A_/Q ^−^ resulting in a slower rate of electron transfer between Q and Q. A reduction in electron transfer, would in turn result in a delay in photosystem II fluorescence induction as it would take longer to reduce the plastoquinone pool and reach maximal fluorescence. To test this hypothesis, we compared the fluorescence induction kinetics of the CA relocalisation lines and wild-type plants. This revealed that there was a 28% increase in the amount of time taken to reach 90% maximal fluorescence in the CA relocalisation lines (Figure 5H). Thus, although change in location of CA activity from the chloroplast to the cytosol did not affect CO_2_ supply to rubisco, the reduction in bicarbonate concentration in the chloroplast impaired the function of photosystem II.

## Discussion

The evolution of carbon concentrating mechanisms in angiosperms, with over 100 independent origins (Gilman et al., 2023; Sage, 2016), is one of the most remarkable examples of convergent evolution in eukaryotic biology. While there is a great diversity in the biochemical and anatomical adaptations that enable these carbon concentrating mechanisms, there are a small number of common features that unify them all. One feature shared by all independent origins of carbon concentrating mechanisms in land plants is the relocalisation of carbonic anhydrase activity from the chloroplast to the cytosol (Ludwig, 2012). Here, we used genome editing to recreate this key evolutionary step in the C_3_ crop rice. We demonstrate that by editing the chloroplast targeting sequence of the primary CA isoform in the leaf that it is possible to change the location of the CA activity from the chloroplast to the cytosol. We show that this change in location of CA activity has no effect on growth or leaf-level CO_2_ assimilation, and uncover a key function for CA in the rapid response of photosystem II to light. Thus, we have revealed a novel role for chloroplast CA in C_3_ plants, and shown that it is possible to recreate a key step in the evolution angiosperm carbon concentrating mechanisms using genome editing.

Several studies have previously investigated the impact of deletion of carbonic anhydrase in C_3_ plants (Hines et al., 2021; Price et al., 1994). A unifying theme of these studies is that removal of CA activity has limited impact on CO_2_ assimilation even when CA is largely reduced or absent (Ferreira et al., 2008; Majeau et al., 1994; Price et al., 1994). The results presented here are consistent with these previous studies, and reveal that the change in location of CA activity from the chloroplast to the cytosol had no effect on leaf level CO_2_ assimilation or growth. However, the discovery that the removal of carbonic anhydrase activity from the chloroplast results in a delay to fluorescence induction provides new insight into the function of chloroplast CA activity in C_3_ plants. It indicates that CA activity is important for facilitating rapid fluorescence induction in the dark to light transition. It does this through maintenance of bicarbonate levels at neutral pH in the dark in the chloroplast to supply bicarbonate to bind to Q_A_.

In this work we have shown that it is possible to achieve a key step in the evolution of C_4_ photosynthesis by genome editing. Previous attempts to recapitulate aspects of C_4_ metabolism or anatomy have all relied on genetic modification (Ermakova et al., 2021; Wang et al., 2017). Given the rapidly changing global regulatory landscape, and the difference in public perception of genetic modification and genome editing (Sprink et al., 2022), it is important to find ways to deliver novel potential yield enhancing technologies through genome editing approaches. While the challenge of delivering a fully operational carbon concentrating mechanism by genome editing is daunting, it may be possible to implement many of the required evolutionary changes by these approaches. For example, many of the key amino acid changes that are required to alter the kinetics of enzymes that function in C_4_ cycle (Niklaus & Kelly, 2019) have the potential to be delivered by genome editing as they require only single nucleotide changes. Moreover, there are now several examples of successful altering of gene expression levels through targeted editing of cis-regulatory sequences (Tang & Zhang, 2023), and thus there is also potential to achieve the upregulation of expression of key enzymes and transporters that are required for engineering of a productive C_4_ cycle. Thus, although the challenge may be daunting, the rapid advances in genome editing technology, coupled with relative ease with which nature has repeatedly evolved the same solution, may mean that it is feasible to using genome editing engineer all or most of a C_4_ cycle in C_3_ plants. In this work we have taken the first step towards engineering a C_4_ cycle in rice by genome editing. Accordingly, this work represents an important first step on the pathway to a genome edited C_4_ rice plant.

## Materials and Methods

### Plant material and growth conditions

All CA-relocalisation lines and wild-type rice (*Oryza sativa*, ssp. japonica, cv. Kitaake) were grown at the University of Oxford in a controlled environment growth chamber. All plants grew under a photoperiod of 14-16 h (8:00 to 22:00 h standard time) with air temperature (*t*_air_) of 26-28 °C, while *t*_air_ in the dark period was of 22 °C. A maximum Photosynthetically active Photon Flux Density (PPFD) was about ∼800-1000 *µ*mol photons m^−2^ s^−1^ for plants grown at Washington State University and ∼300 *µ*mol photons m^−2^ s^−1^ for plants grown in Oxford. Plants were grown in pots in a clay (Oxford) or soil blended with calcined clay in a 4:1 volume ratio (Washington State University).

### Generation of CRISPR constructs

CRISPR constructs were created using MoClo Golden Gate cut-ligations-reactions according to (Hahn et al., 2020) Briefly, 20 bp sgRNA target sites were cut-ligated into level 0 tRNA-sgRNA subcloning vectors via BpiI® (Thermo Fisher) using annealed primer pairs (FHOx6/FHOx7 into pFH51; FHOx8/FHOx9 into pFH51; FHOx10/FHOx11 into pAK002). The generated level 0 tRNA-sgRNA modules were then combined via another cut-ligation reaction using BsaI-HF®v2 (New England Biolabs) into two L1 transcription units (C441001 and C441002) using the rice OsU3 promoter (pFH38), the endlinker pAK-EL-02, the level 1 backbone pICH47751, pAK002+FHOx10/11 and pFH51+FHOx6/FHOx7 or pFH51+FHOx8/FHOx9 respectively. The final vectors C442001 and C442002 were then assembled in a final cut-ligation reaction via BpiI using the respective generated L1 sgRNA transcription units (C441001 or C441002) plus pICSL4723 (L2 backbone), pL1M-R1-pOsAct1-HYG-tNOS-17100 (reversed), pFH66 (ZmUBIp::CAS9:NOSt) and pICH41766 (Endlinker). The vectors pFH38 (Addgene ID #126012), pFH51 (Addgene ID #128213), pAK002 (Addgene ID #128215), pAK-EL-02 (Addgene ID #125761) and pFH66 (Addgene ID #131765) were kindly provided by Vladimir Nekrasov (Rothamsted Research, Harpenden, United Kingdom), pICSL4723 was provided by Mark Youles (Sainsbury Laboratory, Norwich, United Kingdom). pICH47742 (Addgene ID #48001), pICH47751 (Addgene ID #48002) and pICH41766 (Addgene ID #48018) were provided by Sylvestre Marillonnet (Leibniz Institute of Plant Biochemistry, Halle, Germany). pICH41421 (Addgene ID #50339) was provided by Sylvestre Marillonnet and Nicola Patron (Earlham Institute, United Kingdom). All vector maps and sequences generated in this manuscript, as well as all primer sequences, are provided in Supplemental File S5.

### Transformation and recovery of mutant alleles

Rice (cv. Kitaake) calli were transformed using Agrobacterium strain EHA105 with the vectors C442001 or C442002 according to Toki et al., 2006. DNA was isolated from regenerated T_0_ plants (Weigel & Glazebrook, 2009).Presence of the T-DNA was verified by PCR amplification of the *CAS9* gene using DreamTaq®-DNA Polymerase (ThermoFisher Scientific). CRISPR induced mutation were detected by amplification of the *CA1* gene and direct sequencing of the PCR product (Source Biosciences). In some cases, *CA1* PCR products were also subcloned (Zero Blunt™ TOPO™ PCR Cloning Kit, Thermo Fisher) and single clones were sequenced.

Several lines were brought forward into the next generations to segregate out the T-DNA and to recover homozygous *ca1* knockout and frameshift alleles. Absence of T-DNA was verified by DreamTaq® PCR amplification of the *CAS9* gene and the *HPTII* gene as described above and homozygous CRISPR induced *ca1* mutations were verified again by direct sequencing of *ca1* PCR product. Transgene-free, homozygous *ca1* plants were recovered amongst T1 and T2 generation plants, and were used to generate seed stocks for all alleles used in this study.

### Protein localization experiments

Monocot codon optimized coding sequences of the CA1 WT protein, a ca1 protein without target peptide, and the ca1-FS1/2/3/4 protein variants were synthesized as L0 CDS1ns modules according to the MoClo Golden Gate system by Weber et al., 2011 and (Engler et al., 2014) (module names C440011, C440017, C440016, C440013, C440014, C440018, respectively; Gene Universal, United States). Additionally, a C-terminal mTURQUOISE2 L0 CT module was synthesised (C440010). These modules were combined into L1 expression cassettes in a cut-ligation reaction via BsaI-HF®v2 (New England Biolabs) as following: pICH47742, pL0M-PU-pZmUbi-intron1-15455, C440010 and pICH41421 were combined with module C440011 resulting in the L1 vector C441062; or combined with module C440017 resulting in the L1 vector C441068; or with module C440016 resulting in the L1 vector C440067; or with module C440013 resulting in the L1 vector C441064; or with module C440014 resulting in the L1 vector C441065; or with module C440018 resulting in the L1 vector C441069. To create a mTurquoise2 only control, we amplified the mTURQUOISE2 gene from vector C440010 using primers FHOx90/FHOx91 and Q5 DNA Polymerase (New England Biolabs) and combined the PCR product with modules pICH47742, pL0M-PU-pZmUbi-intron1-15455 and pICH41421, resulting in the L1 vector C441070. All L1 and L2 vector maps and sequences as well as all primer sequences are provided in Supplemental File S5.

Protoplasts were extracted from 7 day old rice cv. Kitaake plants following the procedure from (Shan et al., 2014) with the following adaptions: Modifications to the digestion solutions (0.65 M Mannitol, 20 mM MES-KOH, pH 5.7, 10 mM KCl, 5 mM beta-Mercaptoethanol, 1 mM CaCl2, 1 mg/mL BSA, 1.5 % (w/v) Cellulase R-10, 1.5 % (w/v) Cellulase Onozuka RS, 1 % (w/v) Macerozyme R-10, 1 % (w/v) Driselase) allowed us to skip the vacuum infiltration step and proceed directly to a 2h digestion step at 29 °C/70 rpm. After protoplast collection, we added an additional washing step with W5 solution to remove residual enzyme solution. 50,000 protoplasts were transformed with 20 µg of vector DNA (or water control) and fluorescent protein expression was monitored after 2 days of incubation at room temperature using a Zeiss LSM 880 Airy Scan microscope.

### Carbonic Anhydrase activity assay

The assay was performed according to (Hines et al., 2021). Briefly, samples of leaf tissue (250 mg fresh weight) were taken from the longest leaf of 8-week-old rice plants and crushed in liquid nitrogen using a precooled mortar and pestle. A 3:1 (v/w) protein extraction (200mM Tris-HCL, pH 8.3, 1mM EDTA, 20 mM MgCl_2_, 50mM NaCl, 100 mM Na_2_SO_4_) buffer was added to the crushed material to form a homogenate. The homogenate was centrifuged at 4°C for 15 min, and the resulting supernatant was used for the assay. A 200 µL of bromothymol blue solution, comprising 20 mM Tris-HCl, pH 8.3, and 0.2% bromothymol blue, was submitted to a pH-colorimetric reaction by addition of 700 µL of cold, carbonated ddH2O (SodaStream). The time for color-pH change from blue-pH 8.3 to yellow-pH 6.25 was recorded for the uncatalyzed reaction (ts, s), and for the catalyzed reaction (tb, s) in presence of 100 µl of supernatant. The units of CA activity were computed according as (tb-ts)/ts. (Wilbur & Anderson, 1948).

### Chloroplast isolation and CA activity measurement

Chloroplast isolation from rice leaves were performed according to the method described in Flores-Pérez & Jarvis, 2017 with slight modifications. Briefly, rice leaves (10-20g) were washed in 100 mL of cold CIB (0.6 M sorbitol, 10 mM magnesium chloride (MgCl_2_), 10 mM Ethylene Glycol Tetraacetic Acid (EGTA), 10 mM ethylenediaminetetraacetic acid (EDTA), 20 mM sodium bicarbonate (NaHCO_3_), 40 mM 4-(2-hydroxyethyl)piperazine-1-ethanesulfonic acid (HEPES); adjust to pH 8.0 with KOH) on ice and then immediately transferred to 50 ml of cold CIB. The leaves were ground using a blender with short pulses. The homogenate was filtered through a double layer of Miracloth into a 250 mL glass beaker, and the Miracloth was gently squeezed. The extracted chloroplasts were divided into two 50 ml centrifuge tubes and centrifuged at 1000 × g for 5 min at 4 °C. After pouring off the supernatant, the chloroplast pellet was gently resuspended in the remaining supernatant (∼500 μL) by rotating the tube on ice, with additional ice-cold CIB added if necessary. To rinse the chloroplasts, approximately 20–25 mL of HMS buffer (50 mM HEPES, 3 mM magnesium sulfate (MgSO_4_), 0.3 M sorbitol; adjust to pH 8.0 with sodium hydroxide (NaOH)) was added to the tube and inverted carefully. After centrifugation at 1000 × g for 5 min with the brake on, the supernatant was discarded, and the chloroplast pellet was gently resuspended in the remaining HMS buffer, with additional buffer added if required. The chloroplasts were filtered through two layers of Miracloth to remove impurities and then normalized for volume. 100-500 µL of isolated chloroplast was taken in a separate Eppendorf tube and stored in the dark for chlorophyll measurement. 20-500 µL of chloroplast was taken into a separate Eppendorf tube, 1:2 protein extraction buffer was added, vortex vigorously, and centrifuged at 4 °C for 5 min. The supernatant was used to determine CA activity as described previously.

### Chlorophyll fluorescence induction curve measurements

The analysis were done on a LI-6800 portable photosynthetic equipment (LI-COR Biosciences, NE, USA) equipped with a multiphase flash fluorometer (L_ch_; 6800-01A). The measurements were conducted on the detached leaves of 8-weeks old rice plants following a completely randomized design. Leaf temperature (*t*_leaf_) was set at 15 °C, *p*CO_2_ supplied to the leaf chamber (*C*_a_) was 410 μmol CO_2_ mol^−1^ air. Leaf-air VPD (VPD_L_) was in the range of 1.0-1.2 kPa, and air flow through the L_ch_ was 500 *µ*mol air s^−1^. The detached leaves of each plant were allowed to acclimatize in Li-6800 chamber for 20 min in dark to above mentioned conditions followed by 1 s of dark and 8 s of 37 *µ*mol photons m^−2^ s^−1^ red light exposure to find out time for maximum fluorescence. The time taken to reach 90% of F_max_ were calculated from the recorded data.

### Leaf-atmosphere gas exchange: equipment, measurement protocol and functional variables determined

A LI-6800 portable photosynthetic equipment (LI-COR Biosciences, NE, USA) operating as an open system was used, which was equipped with a clear-top chamber (L_ch_; 6800-12A) assembled with a Head Light (LED source: 90% Red; 10 Blue; LI-COR Biosciences, NE, USA). Two O_2_ partial pressures (*p*O_2_, Pa) in the air flow feeding the system were generated by blending different volumetric (∼ molar) fractions of N_2_ and O_2_ through two mass flow controllers (Aalborg, Orangeburg, NY, USA).

The measurements were conducted on the rice cTP-FS and WT plants (n= 4-8) following a completely randomized design. Specifically, for each plant, the mid to distal portions of three fully developed leaves were used to entirely cover a 9 cm^2^ L_ch_ section surface area. *t*_leaf_ was set at 30 °C, *p*CO_2_ supplied to the *C*_a_ was 46.0 Pa (500 *µ*mol CO_2_ mol^−1^ air). VPD_L_ was in the range of 1.0-1.2 kPa, and air flow through the L_ch_ was 350 *µ*mol air s^−1^ (570 mL air min^−1^). In sequence, the measurements were performed at *p*O_2_ of 19.3 kPa (current ambient *p*O_2_; 21% O_2_), then at *p*O_2_ of 1.84 kPa (2% O_2_); PPFD was 1,500 *µ*mol photons m^−2^ s^−1^. The H_2_O molar fraction (mmol H_2_O mol^−1^ air) in the air flow entering the leaf chamber was set and kept constant at each experimental *p*O_2_. Leaf lamina portions were acclimated in the L_ch_ at each *p*O_2_ for circa 30 min; the data were then recorded for 30-40 min. The net CO_2_ assimilation rates per unit (one side) leaf surface area (A, µmol CO_2_ m^−2^ s^−1^), stomatal conductance to CO_2_ diffusion (g_CO2s_, μmol CO_2_ m^−2^ s^−1^ Pa^−1^) and *p*CO_2_ in the intercellular air space (C_i_, Pa) were calculated by LI-6800 software.

### RNA isolation and sequencing

The mid portion of the longest leaf of 8-week-old rice plants was collected and rapidly frozen in liquid nitrogen. Subsequently, the frozen tissue was pulverized into a fine powder using a tissue lyser to facilitate cellular disruption. For each sample, leaf sections from 5 individual plants were combined. Total RNA was extracted using the QIAGEN RNeasy plant mini kit following the manufacturer’s protocol. The concentration and purity of RNA were assessed by measuring absorbance at 260/280 nm and 260/230 nm using a Nanodrop^TM^ spectrophotometer. Three separate sets of RNA from each line were prepared and sent for RNA sequencing (RNA-Seq). Library preparation and RNA sequencing was performed by Novogene. Raw RNA-Seq reads have been deposited to EBI Array express and are available under the accession number E-MTAB-14111. Raw RNA-seq reads were trimmed with fastp (Chen et al., 2018). The *Oryza sativa*, ssp. japonica, cv. Kitaake rice reference transcriptome was downloaded from Phytozome (Goodstein et al., 2011). The gene models for the OsKitaake01g256600 gene in the reference transcriptome and a specific reference transcriptome was also generated for each CA relocalisation line (Supplemental File 2) so that transcript abundance estimates could be accurately computed. Trimmed RNA-seq reads were quantified using the reference transcriptome using Salmon v1.10 (Patro et al., 2017) and differential expression analysis was performed using DESeq 2 (Love et al., 2014). All transcript abundance estimates and difference expression analysis results are provided in Supplemental File 3.

## Acknowledgements

The authors want to thank Chiara Perico for support with microscopy imaging, Julia Lambret Frotte and Daniela Vlad for help with rice transformation, Georgia Smith for help with protoplast isolation protocol set up, Julie Bull, Lizzie Jamison, and Nina Johnson for technical support, and John Baker for photography support. This research was funded by the Bill and Melinda Gates Foundation C_4_ Rice grant awarded to the University of Oxford (2015-2019 (OPP1129902) and 2019-2024 (INV-002970) and the European Union’s Horizon 2020 research and innovation program under grant agreement number 637765. SK was supported by a Royal Society University Research Fellowship.

## Author contributions

SK conceived the project. FH designed all CRIPSR constructs, generated all transgenic plants and conducted the protoplast localisation experiment. JJ performed the growth assay, the carbonic anhydrase assays, the fluorescence induction experiment, and all transcriptomics. NP assisted with the growth and maintenance of plants and in performing CA assays. RG performed leaf gas exchange measurements and photosynthesis analyses. AC contributed to the experimental design and interpretation of the results. SK, FH, and JJ wrote the manuscript. All authors contributed to the editing and revision of the manuscript.

## Data availability

All data is provided in the supplementary information. Raw RNA-Seq reads have been deposited to EBI Array express and are available under the accession number E-MTAB-14111.

## References

Badger, M. (2003). The roles of carbonic anhydrases in photosynthetic CO_2_ concentrating mechanisms. Photosynthesis Research, 77(2), 83–94. 10.1023/A:1025821717773

Badger, M., & Price, G. (1994). The role of carbonic anhydrase in photosynthesis. Annual Review of Plant Biology, 45, 369–392. 10.1146/annurev.pp.45.060194.002101

Brinkert, K., De Causmaecker, S., Krieger-Liszkay, A., Fantuzzi, A., & Rutherford, A. W. (2016). Bicarbonate-induced redox tuning in Photosystem II for regulation and protection. Proceedings of the National Academy of Sciences, 113(43), 12144–12149. 10.1073/pnas.1608862113

Brown, R. H. (1978). A difference in N use efficiency in C_3_ and C_4_ plants and its implications in adaptation and evolution. Crop Science, 18(1), 93–98.

Chen, S., Zhou, Y., Chen, Y., & Gu, J. (2018). fastp: an ultra-fast all-in-one FASTQ preprocessor. Bioinformatics, 34(17), 884–890. 10.1093/bioinformatics/bty560

Clayton, H., Saladié, M., Rolland, V., Sharwood, R., Macfarlane, T., & Ludwig, M. (2017). Loss of the chloroplast transit peptide from an ancestral C_3_ carbonic anhydrase is associated with C_4_ evolution in the grass genus Neurachne. Plant Physiology, 173(3), 1648–1658. 10.1104/pp.16.01893

Cox, N., Jin, L., Jaszewski, A., Smith, P. J., Krausz, E., Rutherford, A. W., & Pace, R. (2009). The semiquinone-iron complex of photosystem II: structural insights from ESR and theoretical simulation; evidence that the native ligand to the non-heme iron is carbonate. Biophysical Journal, 97(7), 2024–2033. 10.1016/j.bpj.2009.06.033

Engler, C., Youles, M., Gruetzner, R., Ehnert, T.-M., Werner, S., Jones, J. D. G., Patron, N. J., & Marillonnet, S. (2014). A golden gate modular cloning toolbox for plants. ACS Synthetic Biology, 3(11), 839–843. 10.1021/sb4001504

Ermakova, M., Arrivault, S., Giuliani, R., Danila, F., Alonso-Cantabrana, H., Vlad, D., Ishihara, H., Feil, R., Guenther, M., Borghi, G. L., Covshoff, S., Ludwig, M., Cousins, A. B., Langdale, J. A., Kelly, S., Lunn, J. E., Stitt, M., von Caemmerer, S., & Furbank, R. T. (2021). Installation of C_4_ photosynthetic pathway enzymes in rice using a single construct. Plant Biotechnol. J., 19(3), 575–588. 10.1111/pbi.13487

Ferreira, F. J., Guo, C., & Coleman, J. R. (2008). Reduction of plastid-localized carbonic anhydrase activity results in reduced Arabidopsis seedling survivorship. Plant Physiology, 147(2), 585–594. 10.1104/pp.108.118661

Flores-Pérez, Ú., & Jarvis, P. (2017). Isolation and suborganellar fractionation of Arabidopsis chloroplasts. Methods Mol Biol, 1511, 45–60. 10.1007/978-1-4939-6533-5_4

Furbank, R. T. (2016). Walking the C_4_ pathway: past, present, and future. Journal of Experimental Botany, 67(14), 4057–4066. 10.1093/jxb/erw161

Gilman, I. S., Smith, J. A. C., Holtum, J. A. M., Sage, R. F., Silvera, K., Winter, K., & Edwards, E. J. (2023). The CAM lineages of planet Earth. Annals of Botany, 132(4), 627–654. 10.1093/aob/mcad135

Goodstein, D. M., Shu, S., Howson, R., Neupane, R., Hayes, R. D., Fazo, J., Mitros, T., Dirks, W., Hellsten, U., Putnam, N., & Rokhsar, D. S. (2011). Phytozome: a comparative platform for green plant genomics. Nucleic Acids Res, 40(Database issue), D1178–86. 10.1093/nar/gkr944

Hahn, F., Korolev, A., Sanjurjo Loures, L., & Nekrasov, V. (2020). A modular cloning toolkit for genome editing in plants. BMC Plant Biology, 20(1), 179. 10.1186/s12870-020-02388-2

Hatch, M. D., & Burnell, J. N. (1990). Carbonic anhydrase activity in leaves and its role in the first step of C_4_ photosynthesis. Plant Physiol, 93(2), 825–828. 10.1104/pp.93.2.825

Hibberd, J. M., Sheehy, J. E., & Langdale, J. A. (2008). Using C_4_ photosynthesis to increase the yield of rice-rationale and feasibility. Curr Opin Plant Biol, 11(2), 228–231. 10.1016/j.pbi.2007.11.002

Hines, K. M., Chaudhari, V., Edgeworth, K. N., Owens, T. G., & Hanson, M. R. (2021). Absence of carbonic anhydrase in chloroplasts affects C_3_ plant development but not photosynthesis. Proceedings of the National Academy of Sciences, 118(33), e2107425118. 10.1073/pnas.2107425118

Lindskog, S., & Coleman, J. E. (1973). The catalytic mechanism of carbonic anhydrase. Proceedings of the National Academy of Sciences, 70(9), 2505–2508. 10.1073/pnas.70.9.2505

Love, M. I., Huber, W., & Anders, S. (2014). Moderated estimation of fold change and dispersion for RNA-seq data with DESeq2. Genome Biology, 15(12), 550. 10.1186/s13059-014-0550-8

Ludwig, M. (2012). Carbonic anhydrase and the molecular evolution of C_4_ photosynthesis. Plant, Cell & Environment, 35(1), 22–37. 10.1111/j.1365-3040.2011.02364.x

Ludwig, M. (2016). Evolution of carbonic anhydrase in C_4_ plants. Current Opinion in Plant Biology, 31, 16–22. 10.1016/j.pbi.2016.03.003

Majeau, N., Arnoldo, M. A., & Coleman, J. R. (1994). Modification of carbonic anhydrase activity by antisense and over-expression constructs in transgenic tobacco. Plant Mol Biol, 25(3), 377–385. 10.1007/BF00043867

Mattinson, O., & Kelly, S. (2024). The metabolite transporters of C_4_ photosynthesis. 10.32942/x2f89t

Murray, A., Mendieta, J. P., Vollmers, C., & Schmitz, R. J. (2022). Simple and accurate transcriptional start site identification using Smar2C2 and examination of conserved promoter features. The Plant Journal, 112(2), 583–596. 10.1111/tpj.15957

Niklaus, M., & Kelly, S. (2019). The molecular evolution of C_4_ photosynthesis: opportunities for understanding and improving the world’s most productive plants. Journal of Experimental Botany, 70(3), 795–804. 10.1093/jxb/ery416

Okabe, K., Yang, S.-Y., Tsuzuki, M., & Miyachi, S. (1984). Carbonic anhydrase: Its content in spinach leaves and its taxonomic diversity studied with anti-spinach leaf carbonic anhydrase antibody. Plant Science Letters, 33(2), 145–153. 10.1016/0304-4211(84)90004-X

Patro, R., Duggal, G., Love, M. I., Irizarry, R. A., & Kingsford, C. (2017). Salmon provides fast and bias-aware quantification of transcript expression. Nat Methods, 14(4), 417–419. 10.1038/nmeth.4197

Poincelot, R. P. (1972). The distribution of carbonic anhydrase and ribulose diphosphate carboxylase in maize leaves. Plant Physiol, 50(3), 336–340. 10.1104/pp.50.3.336

Price, G. D., von Caemmerer, S., Evans, J. R., Yu, J.-W., Lloyd, J., Oja, V., Kell, P., Harrison, K., Gallagher, A., & Badger, M. R. (1994). Specific reduction of chloroplast carbonic anhydrase activity by antisense RNA in transgenic tobacco plants has a minor effect on photosynthetic CO2 assimilation. Planta, 193(3), 331–340. 10.1007/BF00201810

Sage, R. F. (2016). A portrait of the C_4_ photosynthetic family on the 50th anniversary of its discovery: species number, evolutionary lineages, and Hall of Fame. Journal of Experimental Botany, 67(14), 4039–4056. 10.1093/jxb/erw156

Shan, Q., Wang, Y., Li, J., & Gao, C. (2014). Genome editing in rice and wheat using the CRISPR/Cas system. Nature Protocols, 9(10), 2395–2410. 10.1038/nprot.2014.157

Sprink, T., Wilhelm, R., & Hartung, F. (2022). Genome editing around the globe: An update on policies and perceptions. Plant Physiol, 190(3), 1579–1587. 10.1093/plphys/kiac359

Tang, X., & Zhang, Y. (2023). Beyond knockouts: fine-tuning regulation of gene expression in plants with CRISPR-Cas-based promoter editing. New Phytologist, 239(3), 868–874. 10.1111/nph.19020

Tanz, S. K., Tetu, S. G., Vella, N. G. F., & Ludwig, M. (2009). Loss of the transit peptide and an increase in gene expression of an ancestral chloroplastic carbonic anhydrase were instrumental in the evolution of the cytosolic C_4_ carbonic anhydrase in Flaveria. Plant Physiology, 150(3), 1515–1529. 10.1104/pp.109.137513

Toki, S., Hara, N., Ono, K., Onodera, H., Tagiri, A., Oka, S., & Tanaka, H. (2006). Early infection of scutellum tissue with Agrobacterium allows high-speed transformation of rice. Plant J, 47(6), 969–976. 10.1111/j.1365-313X.2006.02836.x

Tsuzuki, M., Miyachi, S., & Edwards, G. E. (1985). Localization of carbonic anhydrase in mesophyll cells of terrestrial C_3_ plants in relation to CO_2_ assimilation. Plant and Cell Physiology, 26(5), 881–891. 10.1093/oxfordjournals.pcp.a076983

von Caemmerer, S., Quick, W. P., & Furbank, R. T. (2012). The development of C₄ rice: current progress and future challenges. Science, 336(6089), 1671–1672. 10.1126/science.1220177

Walker, B., VanLoocke, A., Bernacchi, C., & Ort, D. (2016). The Costs of Photorespiration to Food Production Now and in the Future. Annual Review of Plant Biology, 67. 10.1146/annurev-arplant-043015-111709

Wang, P., Khoshravesh, R., Karki, S., Tapia, R., Balahadia, C. P., Bandyopadhyay, A., Quick, W. P., Furbank, R., Sage, T. L., & Langdale, J. A. (2017). Re-creation of a Key Step in the Evolutionary Switch from C_3_ to C_4_ Leaf Anatomy. Curr Biol, 27(21), 3278–3287.e6. 10.1016/j.cub.2017.09.040

Way, D. A., Katul, G. G., Manzoni, S., & Vico, G. (2014). Increasing water use efficiency along the C_3_ to C_4_ evolutionary pathway: a stomatal optimization perspective. Journal of Experimental Botany, 65(13), 3683–3693. 10.1093/jxb/eru205

Weber, E., Engler, C., Gruetzner, R., Werner, S., & Marillonnet, S. (2011). A Modular cloning system for standardized assembly of multigene constructs. PLOS ONE, 6(2), e16765-. 10.1371/journal.pone.0016765

Weigel, D., & Glazebrook, J. (2009). Quick miniprep for plant DNA isolation. Cold Spring Harb Protoc, 2009(3), db.prot5179. 10.1101/pdb.prot5179

Wilbur, K. M., & Anderson, N. G. (1948). Electrometric and colorimetric determination of carbonic anhydrase. J Biol Chem, 176(1), 147–154. https://www.ncbi.nlm.nih.gov/pubmed/18886152

Zhu, X.-G., Long, S. P., & Ort, D. R. (2008). What is the maximum efficiency with which photosynthesis can convert solar energy into biomass? Current Opinion in Biotechnology, 19(2), 153–159. 10.1016/j.copbio.2008.02.004

