## Supplemental File 1 for "Realisation of a key step in the evolution of C_4_ photosynthesis in rice by genome editing"

### Figure S1

| **CA Type** | **O. sativa** | **O. sativa Kitaake** | **WT1** | **WT2** | **WT3** | **WT4** | **WT5** |
| --- | --- | --- | --- | --- | --- | --- | --- |
| Alpha | LOC_Os04g33660.2 | OsKitaake04g120500 | 107 | 105 | 114 | 102 | 95 |
| Alpha | LOC_Os02g33030.1 | OsKitaake02g195200 | 7 | 7 | 9 | 10 | 9 |
| Alpha | LOC_Os06g40770.1 | OsKitaake06g211400 | 0 | 0 | 0 | 0 | 0 |
| Alpha | LOC_Os08g32750.1 | OsKitaake08g156600 | 0 | 0 | 0 | 0 | 0 |
| Alpha | LOC_Os08g32780.1 | OsKitaake08g156700 | 0 | 0 | 0 | 0 | 0 |
| Alpha | LOC_Os08g32840.1 | OsKitaake08g156800 | 0 | 0 | 0 | 0 | 0 |
| Alpha | LOC_Os08g36630.1 | OsKitaake08g183100 | 0 | 0 | 0 | 0 | 0 |
| Alpha | LOC_Os08g36680.1 | OsKitaake08g183200 | 0 | 0 | 0 | 0 | 0 |
| Alpha | LOC_Os09g28150.1 | OsKitaake09g124500 | 0 | 0 | 0 | 0 | 0 |
| Alpha | LOC_Os11g05510.1 | OsKitaake11g035500 | 0 | 0 | 0 | 0 | 0 |
| Alpha | LOC_Os12g05730.1 | OsKitaake12g040400 | 0 | 0 | 0 | 0 | 0 |
| **Beta** | **LOC_Os01g45274** | **OsKitaake01g256600** | **16418** | **16110** | **16085** | **17241** | **17843** |
| Beta | LOC_Os09g28910 | OsKitaake09g130100 | 40 | 41 | 38 | 39 | 37 |
| Gamma | LOC_Os01g18070.1 | OsKitaake01g129000 | 54 | 54 | 53 | 55 | 49 |
| Gamma | LOC_Os12g07220.1 | OsKitaake12g050700 | 21 | 20 | 19 | 22 | 17 |
| Gamma | LOC_Os07g44840.1 | OsKitaake07g248500 | 1 | 0 | 1 | 1 | 1 |
| Gamma-Like | LOC_Os02g30460.1 | OsKitaake02g182500 | 11 | 13 | 13 | 13 | 12 |

**Figure S1.** mRNA abundance of all carbonic anhydrase isoforms in fully expanded leaves of rice. The carbonic anhydrase type is provided as well as the accession numbers in both the *Oryza sativa ssp. japonica* genome reference and the genome reference for *Oryza sativa ssp. japonica* cv. KitaakeX. mRNA abundance values provided are in transcripts per million. Five pooled biological replicates of wild type (WT) leaves are shown. The gene encoding the chloroplast localize beta carbonic anhydrase is highlighted in bold green font. Data from Danila *et. al* Plant Biotechnology Journal. 2022 20(9):1786-1806.

### Figure S2

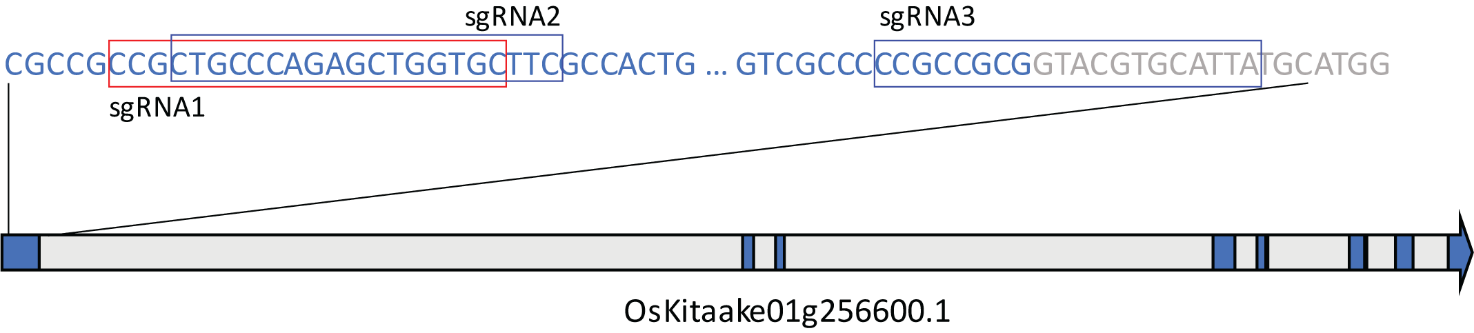

**Figure S2**. Gene model and guide RNA targets sequences. Exons are shown in blue and introns shown in grey. Exon sequence is shown in blue font and intron sequence shown in grey font. The three different guide RNA sequences are indicated by different colour boxes.

### Figure S3

| **Gene** | **Prediction** | **noTP** | **SP** | **mTP** | **cTP** | **luTP** |
| --- | --- | --- | --- | --- | --- | --- |
| OsKitaake01g256600_T1 | cTP | 0.00 | 0.00 | 0.00 | **1.00** | 0.00 |
| OsKitaake01g256600_T2 | noTP | **1.00** | 0.00 | 0.00 | 0.00 | 0.00 |
| OsKitaake01g256600_T3 | noTP | **1.00** | 0.00 | 0.00 | 0.00 | 0.00 |
| cTP-FS1 | cTP | 0.21 | 0.01 | 0.10 | **0.54** | 0.14 |
| cTP-FS2 | noTP | **0.74** | 0.02 | 0.00 | 0.18 | 0.06 |
| cTP-FS3 | noTP | **0.73** | 0.09 | 0.00 | 0.10 | 0.08 |
| cTP-FS4 | cTP | 0.27 | 0.01 | 0.00 | **0.72** | 0.00 |

**Figure S3.** TargetP subcellular target predictions for wild type and cTP-FS frame shift alleles. Prediction values are shown for the three wild-type transcript variants from the OsKitaake01g256600 gene as well as all four genome edited cTP-FS alleles. The cTP-KO alleles are not shown as these are all short truncated proteins that do not encode a carbonic anhydrase domain.
