## Supplemental File 2 for "Realisation of a key step in the evolution of C_4_ photosynthesis in rice by genome editing"

### Mutant allele sequences in the genome edited rice lines

#### Nucleotide sequences of cTP-FS alleles

##### Legend

Alternating grey and no-shading indicate successive exons

##### X = inserted nucleotides by genome editing

~~ACGT~~ = deleted nucleotides by genome editing

X = Substitution

##### Sequences

>WT_CA1_OsKitaake01g256600.1

ATGTCGACCGCCGCCGCCGCCGCCGCTGCCCAGAGCTGGTGCTTCGCCACTGTCACCCCGCGCTCCCGCGCCACAGTCGTCGCCAGCCTCGCCTCCCCATCACCGTCCTCCTCCTCCTCCTCCTCCAACAGCAGCAACCTCCCGGCCCCCTTCCGCCCCCGCCTCATCCGCAACACCCCCGTCTTCGCCGCCCCCGTCGCCCCCGCCGCGATGGACGCCGCCGTCGACCGCCTCAAGGATGGGTTCGCCAAGTTCAAGACCGAGTTCTATGACAAGAAGCCGGAGCTCTTCGAGCCGCTCAAGGCCGGCCAGGCACCCAAGTACATGGTGTTCTCGTGCGCCGACTCTCGCGTGTGCCCGTCGGTGACCATGGGCCTGGAGCCCGGCGAGGCCTTCACCGTCCGCAACATCGCCAACATGGTCCCAGCTTACTGCAAGATCAAGCACGCTGGCGTCGGGTCGGCCATCGAGTACGCCGTCTGCGCCCTCAAGGTCGAACTCATCGTGGTGATTGGCCACAGCCGCTGCGGTGGAATCAAGGCCCTCCTCTCACTCAAGGATGGAGCACCAGACTCCTTCCACTTCGTCGAGGACTGGGTCAGGACCGGTTTCCCCGCCAAGAAGAAGGTTCAGACCGAGCACGCCTCGCTGCCTTTCGATGACCAATGCGCCATCTTGGAGAAGGAGGCCGTGAACCAATCCCTGGAGAACCTCAAGACCTACCCGTTCGTCAAGGAGGGGATCGCCAACGGCACCCTCAAGCTCGTCGGCGGCCACTACGACTTCGTCTCCGGCAACTTGGACTTATGGGAGCCCTAA

>cTP-FS1

ATGTCGACCGCCGCCGCCGCCGCCGCT**A**~~CCCAG~~AGCTGGTGCTTCGCCACTGTCACCCCGCGCTCCCGCGCCACAGTCGTCGCCAGCCTCGCCTCCCCATCACCGTCCTCCTCCTCCTCCTCCTCCAACAGCAGCAACCTCCCGGCCCCCTTCCGCCCCCGCCTCATCCGCAACACCCCCGTCTTCGCCGCCCCCGTCGCCCCC~~G~~CCGCGATGGACGCCGCCGTCGACCGCCTCAAGGATGGGTTCGCCAAGTTCAAGACCGAGTTCTATGACAAGAAGCCGGAGCTCTTCGAGCCGCTCAAGGCCGGCCAGGCACCCAAGTACATGGTGTTCTCGTGCGCCGACTCTCGCGTGTGCCCGTCGGTGACCATGGGCCTGGAGCCCGGCGAGGCCTTCACCGTCCGCAACATCGCCAACATGGTCCCAGCTTACTGCAAGATCAAGCACGCTGGCGTCGGGTCGGCCATCGAGTACGCCGTCTGCGCCCTCAAGGTCGAACTCATCGTGGTGATTGGCCACAGCCGCTGCGGTGGAATCAAGGCCCTCCTCTCACTCAAGGATGGAGCACCAGACTCCTTCCACTTCGTCGAGGACTGGGTCAGGACCGGTTTCCCCGCCAAGAAGAAGGTTCAGACCGAGCACGCCTCGCTGCCTTTCGATGACCAATGCGCCATCTTGGAGAAGGAGGCCGTGAACCAATCCCTGGAGAACCTCAAGACCTACCCGTTCGTCAAGGAGGGGATCGCCAACGGCACCCTCAAGCTCGTCGGCGGCCACTACGACTTCGTCTCCGGCAACTTGGACTTATGGGAGCCCTAA

>cTP-FS2

ATGTCGACCGCCGCCGCCGCCGCCG**A**CTGCCCAGAGCTGGTGCTTCGCCACTGTCACCCCGCGCTCCCGCGCCACAGTCGTCGCCAGCCTCGCCTCCCCATCACCGTCCTCCTCCTCCTCCTCCTCCAACAGCAGCAACCTCCCGGCCCCCTTCCGCCCCCGCCTCATCCGCAACACCCCCGTCTTCGCCGCCCCCGTCGCCCCCG**AC**CCGCGATGGACGCCGCCGTCGACCGCCTCAAGGATGGGTTCGCCAAGTTCAAGACCGAGTTCTATGACAAGAAGCCGGAGCTCTTCGAGCCGCTCAAGGCCGGCCAGGCACCCAAGTACATGGTGTTCTCGTGCGCCGACTCTCGCGTGTGCCCGTCGGTGACCATGGGCCTGGAGCCCGGCGAGGCCTTCACCGTCCGCAACATCGCCAACATGGTCCCAGCTTACTGCAAGATCAAGCACGCTGGCGTCGGGTCGGCCATCGAGTACGCCGTCTGCGCCCTCAAGGTCGAACTCATCGTGGTGATTGGCCACAGCCGCTGCGGTGGAATCAAGGCCCTCCTCTCACTCAAGGATGGAGCACCAGACTCCTTCCACTTCGTCGAGGACTGGGTCAGGACCGGTTTCCCCGCCAAGAAGAAGGTTCAGACCGAGCACGCCTCGCTGCCTTTCGATGACCAATGCGCCATCTTGGAGAAGGAGGCCGTGAACCAATCCCTGGAGAACCTCAAGACCTACCCGTTCGTCAAGGAGGGGATCGCCAACGGCACCCTCAAGCTCGTCGGCGGCCACTACGACTTCGTCTCCGGCAACTTGGACTTATGGGAGCCCTAA

>cTP-FS3

ATGTCGACCGCCGCCGCCGCCGCCG**A**CTGCCCAGAGCTGGTGCTTCGCCACTGTCACCCCGCGCTCCCGCGCCACAGTCGTCGCCAGCCTCGCCTCCCCATCACCGTCCTCCTCCTCCTCCTCCTCCAACAGCAGCAACCTCCCGGCCCCCTTCCGCCCCCGCC**C**CATCCGCAACACC~~CCCGTCTTCGCCGCCCCCGTCGCCCCCG~~CCGCGATGGACGCCGCCGTCGACCGCCTCAAGGATGGGTTCGCCAAGTTCAAGACCGAGTTCTATGACAAGAAGCCGGAGCTCTTCGAGCCGCTCAAGGCCGGCCAGGCACCCAAGTACATGGTGTTCTCGTGCGCCGACTCTCGCGTGTGCCCGTCGGTGACCATGGGCCTGGAGCCCGGCGAGGCCTTCACCGTCCGCAACATCGCCAACATGGTCCCAGCTTACTGCAAGATCAAGCACGCTGGCGTCGGGTCGGCCATCGAGTACGCCGTCTGCGCCCTCAAGGTCGAACTCATCGTGGTGATTGGCCACAGCCGCTGCGGTGGAATCAAGGCCCTCCTCTCACTCAAGGATGGAGCACCAGACTCCTTCCACTTCGTCGAGGACTGGGTCAGGACCGGTTTCCCCGCCAAGAAGAAGGTTCAGACCGAGCACGCCTCGCTGCCTTTCGATGACCAATGCGCCATCTTGGAGAAGGAGGCCGTGAACCAATCCCTGGAGAACCTCAAGACCTACCCGTTCGTCAAGGAGGGGATCGCCAACGGCACCCTCAAGCTCGTCGGCGGCCACTACGACTTCGTCTCCGGCAACTTGGACTTATGGGAGCCCTAA

>cTP-FS4

ATGTCGACCGCCGCCGCC~~GCCGCCG~~CTGCCCAGAGCTGGTGCTTCGCCACTGTCACCCCGCGCTCCCGCGCCACAGTCGTCGCCAGCCTCGCCTCCCCATCACCGTCCTCCTCCTCCTCCTCCTCCAACAGCAGCAACCTCCCGGCCCCCTTCCGCCCCCGCCTCATCCGCAACACCCCCGTCTTCGCCGCCCCCGTCGCCCCCG**T**CCGCGATGGACGCCGCCGTCGACCGCCTCAAGGATGGGTTCGCCAAGTTCAAGACCGAGTTCTATGACAAGAAGCCGGAGCTCTTCGAGCCGCTCAAGGCCGGCCAGGCACCCAAGTACATGGTGTTCTCGTGCGCCGACTCTCGCGTGTGCCCGTCGGTGACCATGGGCCTGGAGCCCGGCGAGGCCTTCACCGTCCGCAACATCGCCAACATGGTCCCAGCTTACTGCAAGATCAAGCACGCTGGCGTCGGGTCGGCCATCGAGTACGCCGTCTGCGCCCTCAAGGTCGAACTCATCGTGGTGATTGGCCACAGCCGCTGCGGTGGAATCAAGGCCCTCCTCTCACTCAAGGATGGAGCACCAGACTCCTTCCACTTCGTCGAGGACTGGGTCAGGACCGGTTTCCCCGCCAAGAAGAAGGTTCAGACCGAGCACGCCTCGCTGCCTTTCGATGACCAATGCGCCATCTTGGAGAAGGAGGCCGTGAACCAATCCCTGGAGAACCTCAAGACCTACCCGTTCGTCAAGGAGGGGATCGCCAACGGCACCCTCAAGCTCGTCGGCGGCCACTACGACTTCGTCTCCGGCAACTTGGACTTATGGGAGCCCTAA

#### Amino acid sequences of cTP-FS alleles

##### Legend

Chloroplast Transit Peptide

Carbonic anhydrase domain PF00484

Out of frame region

Chloroplast transit peptide cleavage Site (**X^X**)

##### Sequences

>WT_CA1_OsKitaake01g256600.1.p

MSTAAAAAAAQSWCFATVTPRSRATVVASLASPSPSSSSSSSNSSNLPAPFRPRLIRNTPV**F^A**APVAPAAMDAAVDRLKDGFAKFKTEFYDKKPELFEPLKAGQAPKYMVFSCADSRVCPSVTMGLEPGEAFTVRNIANMVPAYCKIKHAGVGSAIEYAVCALKVELIVVIGHSRCGGIKALLSLKDGAPDSFHFVEDWVRTGFPAKKKVQTEHASLPFDDQCAILEKEAVNQSLENLKTYPFVKEGIANGTLKLVGGHYDFVSGNLDLWEP

>cTP-FS1

MSTAAAAAAKLVLRHCHPALPRHSRRQPRLPITVLLLLLLQQQQPPGPLPPPPHPQHPRLRRPRRPPAMDAAVDRLKDGFAKFKTEFYDKKPELFEPLKAGQAPKYMVFSCADSRVCPSVTMGLEPGEAFTVRNIANMVPAYCKIKHAGVGSAIEYAVCALKVELIVVIGHSRCGGIKALLSLKDGAPDSFHFVEDWVRTGFPAKKKVQTEHASLPFDDQCAILEKEAVNQSLENLKTYPFVKEGIANGTLKLVGGHYDFVSGNLDLWEP

>cTP-FS2

MSTAAAAADCPELVLRHCHPALPRHSRRQPRLPITVLLLLLLQQQQPPGPLPPPPHPQHPRLRRPRRPRPAMDAAVDRLKDGFAKFKTEFYDKKPELFEPLKAGQAPKYMVFSCADSRVCPSVTMGLEPGEAFTVRNIANMVPAYCKIKHAGVGSAIEYAVCALKVELIVVIGHSRCGGIKALLSLKDGAPDSFHFVEDWVRTGFPAKKKVQTEHASLPFDDQCAILEKEAVNQSLENLKTYPFVKEGIANGTLKLVGGHYDFVSGNLDLWEP

>cTP-FS3

MSTAAAAADCPELVLRHCHPALPRHSRRQPRLPITVLLLLLLQQQQPPGPLPPPPHPQHPAMDAAVDRLKDGFAKFKTEFYDKKPELFEPLKAGQAPKYMVFSCADSRVCPSVTMGLEPGEAFTVRNIANMVPAYCKIKHAGVGSAIEYAVCALKVELIVVIGHSRCGGIKALLSLKDGAPDSFHFVEDWVRTGFPAKKKVQTEHASLPFDDQCAILEKEAVNQSLENLKTYPFVKEGIANGTLKLVGGHYDFVSGNLDLWEP

>cTP-FS4

MSTAAALPRAGASPLSPRAPAPQSSPASPPHHRPPPPPPPTAATSRPPSAPASSATPPSSPPPSPPSAMDAAVDRLKDGFAKFKTEFYDKKPELFEPLKAGQAPKYMVFSCADSRVCPSVTMGLEPGEAFTVRNIANMVPAYCKIKHAGVGSAIEYAVCALKVELIVVIGHSRCGGIKALLSLKDGAPDSFHFVEDWVRTGFPAKKKVQTEHASLPFDDQCAILEKEAVNQSLENLKTYPFVKEGIANGTLKLVGGHYDFVSGNLDLWEP

#### Nucleotide sequences of cTP-KO alleles

##### Legend

Alternating grey and no-shading indicate successive exons

##### X = inserted nucleotides by genome editing

~~ACGT~~ = deleted nucleotides by genome editing

##### Sequences

>WT_CA1_OsKitaake01g256600.1

ATGTCGACCGCCGCCGCCGCCGCCGCTGCCCAGAGCTGGTGCTTCGCCACTGTCACCCCGCGCTCCCGCGCCACAGTCGTCGCCAGCCTCGCCTCCCCATCACCGTCCTCCTCCTCCTCCTCCTCCAACAGCAGCAACCTCCCGGCCCCCTTCCGCCCCCGCCTCATCCGCAACACCCCCGTCTTCGCCGCCCCCGTCGCCCCCGCCGCGATGGACGCCGCCGTCGACCGCCTCAAGGATGGGTTCGCCAAGTTCAAGACCGAGTTCTATGACAAGAAGCCGGAGCTCTTCGAGCCGCTCAAGGCCGGCCAGGCACCCAAGTACATGGTGTTCTCGTGCGCCGACTCTCGCGTGTGCCCGTCGGTGACCATGGGCCTGGAGCCCGGCGAGGCCTTCACCGTCCGCAACATCGCCAACATGGTCCCAGCTTACTGCAAGATCAAGCACGCTGGCGTCGGGTCGGCCATCGAGTACGCCGTCTGCGCCCTCAAGGTCGAACTCATCGTGGTGATTGGCCACAGCCGCTGCGGTGGAATCAAGGCCCTCCTCTCACTCAAGGATGGAGCACCAGACTCCTTCCACTTCGTCGAGGACTGGGTCAGGACCGGTTTCCCCGCCAAGAAGAAGGTTCAGACCGAGCACGCCTCGCTGCCTTTCGATGACCAATGCGCCATCTTGGAGAAGGAGGCCGTGAACCAATCCCTGGAGAACCTCAAGACCTACCCGTTCGTCAAGGAGGGGATCGCCAACGGCACCCTCAAGCTCGTCGGCGGCCACTACGACTTCGTCTCCGGCAACTTGGACTTATGGGAGCCCTAA

>cTP-KO1

ATGTCGACCGCCGCCGCCGCCGCCGCTGCCCAGAGCTGGTGCTTCGCCACTGTCACCCCGCGCTCCCGCGCCACAGTCGTCGCCAGCCTCGCCTCCCCATCACCGTCCTCCTCCTCCTCCTCCTCCAACAGCAGCAACCTCCCGGCCCCCTTCCGCCCCCGCCTCATCCGCAACACCCCCGTCTTCGCCGCCCCCGTCGCCCCCG**A**CCGCGATGGACGCCGCCGTCGACCGCCTCAAGGATGGGTTCGCCAAGTTCAAGACCGAGTTCTATGACAAGAAGCCGGAGCTCTTCGAGCCGCTCAAGGCCGGCCAGGCACCCAAGTACATGGTGTTCTCGTGCGCCGACTCTCGCGTGTGCCCGTCGGTGACCATGGGCCTGGAGCCCGGCGAGGCCTTCACCGTCCGCAACATCGCCAACATGGTCCCAGCTTACTGCAAGATCAAGCACGCTGGCGTCGGGTCGGCCATCGAGTACGCCGTCTGCGCCCTCAAGGTCGAACTCATCGTGGTGATTGGCCACAGCCGCTGCGGTGGAATCAAGGCCCTCCTCTCACTCAAGGATGGAGCACCAGACTCCTTCCACTTCGTCGAGGACTGGGTCAGGACCGGTTTCCCCGCCAAGAAGAAGGTTCAGACCGAGCACGCCTCGCTGCCTTTCGATGACCAATGCGCCATCTTGGAGAAGGAGGCCGTGAACCAATCCCTGGAGAACCTCAAGACCTACCCGTTCGTCAAGGAGGGGATCGCCAACGGCACCCTCAAGCTCGTCGGCGGCCACTACGACTTCGTCTCCGGCAACTTGGACTTATGGGAGCCCTAA

>cTP-KO2

ATGTCGACCGCCGCCGCCGCCGCCGCTGCCCAGAGCTGGTGCTTCGCCACTGTCACCCCGCGCTCCCGCGCCACAGTCGTCGCCAGCCTCGCCTCCCCATCACCGTCCTCCTCCTCCTCCTCCTCCAACAGCAGCAACCTCCCGGCCCCCTTCCGCCCCCGCCTCATCCGCAACACCCCCGTCTTCGCCGCCCCCGTCGCCCCCG**T**CCGCGATGGACGCCGCCGTCGACCGCCTCAAGGATGGGTTCGCCAAGTTCAAGACCGAGTTCTATGACAAGAAGCCGGAGCTCTTCGAGCCGCTCAAGGCCGGCCAGGCACCCAAGTACATGGTGTTCTCGTGCGCCGACTCTCGCGTGTGCCCGTCGGTGACCATGGGCCTGGAGCCCGGCGAGGCCTTCACCGTCCGCAACATCGCCAACATGGTCCCAGCTTACTGCAAGATCAAGCACGCTGGCGTCGGGTCGGCCATCGAGTACGCCGTCTGCGCCCTCAAGGTCGAACTCATCGTGGTGATTGGCCACAGCCGCTGCGGTGGAATCAAGGCCCTCCTCTCACTCAAGGATGGAGCACCAGACTCCTTCCACTTCGTCGAGGACTGGGTCAGGACCGGTTTCCCCGCCAAGAAGAAGGTTCAGACCGAGCACGCCTCGCTGCCTTTCGATGACCAATGCGCCATCTTGGAGAAGGAGGCCGTGAACCAATCCCTGGAGAACCTCAAGACCTACCCGTTCGTCAAGGAGGGGATCGCCAACGGCACCCTCAAGCTCGTCGGCGGCCACTACGACTTCGTCTCCGGCAACTTGGACTTATGGGAGCCCTAA

>cTP-KO3

ATGTCGACCGCCGCCGCCGCCGCCG**A**~~CTGCCCAGAGCTGGTGCTTCGCCACTGTCACCCCGCGCTCCCGCGCCACAGTCGTCGCCAGCCTCGCCTCCCCATCACCGTCCTCCTCCTCCTCCTCCTCCAACAGCAGCAACCTCCCGGCCCCCTTCCGCCCCCGCCTCATCCGCAACACCCCCGTCTTCGCCGCCCCCGTCGCCCCCG~~CCGCGATGGACGCCGCCGTCGACCGCCTCAAGGATGGGTTCGCCAAGTTCAAGACCGAGTTCTATGACAAGAAGCCGGAGCTCTTCGAGCCGCTCAAGGCCGGCCAGGCACCCAAGTACATGGTGTTCTCGTGCGCCGACTCTCGCGTGTGCCCGTCGGTGACCATGGGCCTGGAGCCCGGCGAGGCCTTCACCGTCCGCAACATCGCCAACATGGTCCCAGCTTACTGCAAGATCAAGCACGCTGGCGTCGGGTCGGCCATCGAGTACGCCGTCTGCGCCCTCAAGGTCGAACTCATCGTGGTGATTGGCCACAGCCGCTGCGGTGGAATCAAGGCCCTCCTCTCACTCAAGGATGGAGCACCAGACTCCTTCCACTTCGTCGAGGACTGGGTCAGGACCGGTTTCCCCGCCAAGAAGAAGGTTCAGACCGAGCACGCCTCGCTGCCTTTCGATGACCAATGCGCCATCTTGGAGAAGGAGGCCGTGAACCAATCCCTGGAGAACCTCAAGACCTACCCGTTCGTCAAGGAGGGGATCGCCAACGGCACCCTCAAGCTCGTCGGCGGCCACTACGACTTCGTCTCCGGCAACTTGGACTTATGGGAGCCCTAA

>cTP-KO4

ATGTCGACCGCCGCCGCCGCCGCCG**T**CTGCCCAGAGCTGGTGCTTCGCCACTGTCACCCCGCGCTCCCGCGCCACAGTCGTCGCCAGCCTCGCCTCCCCATCACCGTCCTCCTCCTCCTCCTCCTCCAACAGCAGCAACCTCCCGGCCCCCTTCCGCCCCCGCCTCATCCGCAACACCCCCGTCTTCGCCGCCCCCGTCGCCCCCG**T**CCGCGATGGACGCCGCCGTCGACCGCCTCAAGGATGGGTTCGCCAAGTTCAAGACCGAGTTCTATGACAAGAAGCCGGAGCTCTTCGAGCCGCTCAAGGCCGGCCAGGCACCCAAGTACATGGTGTTCTCGTGCGCCGACTCTCGCGTGTGCCCGTCGGTGACCATGGGCCTGGAGCCCGGCGAGGCCTTCACCGTCCGCAACATCGCCAACATGGTCCCAGCTTACTGCAAGATCAAGCACGCTGGCGTCGGGTCGGCCATCGAGTACGCCGTCTGCGCCCTCAAGGTCGAACTCATCGTGGTGATTGGCCACAGCCGCTGCGGTGGAATCAAGGCCCTCCTCTCACTCAAGGATGGAGCACCAGACTCCTTCCACTTCGTCGAGGACTGGGTCAGGACCGGTTTCCCCGCCAAGAAGAAGGTTCAGACCGAGCACGCCTCGCTGCCTTTCGATGACCAATGCGCCATCTTGGAGAAGGAGGCCGTGAACCAATCCCTGGAGAACCTCAAGACCTACCCGTTCGTCAAGGAGGGGATCGCCAACGGCACCCTCAAGCTCGTCGGCGGCCACTACGACTTCGTCTCCGGCAACTTGGACTTATGGGAGCCCTAA

#### Amino acid sequences of cTP-KO alleles

##### Legend

Chloroplast Transit Peptide

Carbonic anhydrase domain PF00484

Out of frame region

***** - premature stop codon

Chloroplast transit peptide cleavage Site (**X^X**)

##### Sequences

>WT_CA1_OsKitaake01g256600.1.p

MSTAAAAAAAQSWCFATVTPRSRATVVASLASPSPSSSSSSSNSSNLPAPFRPRLIRNTPV**F^A**APVAPAAMDAAVDRLKDGFAKFKTEFYDKKPELFEPLKAGQAPKYMVFSCADSRVCPSVTMGLEPGEAFTVRNIANMVPAYCKIKHAGVGSAIEYAVCALKVELIVVIGHSRCGGIKALLSLKDGAPDSFHFVEDWVRTGFPAKKKVQTEHASLPFDDQCAILEKEAVNQSLENLKTYPFVKEGIANGTLKLVGGHYDFVSGNLDLWEP

>cTP-KO1

MSTAAAAAAAQSWCFATVTPRSRATVVASLASPSPSSSSSSSNSSNLPAPFRPRLIRNTPV**F^A**APVAPDRDGRRRRPPQGWVRQVQDRVL*****

>cTP-KO2 MSTAAAAAAAQSWCFATVTPRSRATVVASLASPSPSSSSSSSNSSNLPAPFRPRLIRNTPV**F^A**APVAPVRDGRRRRPPQGWVRQVQDRVL*

>cTP-KO3

MSTAAAAADRDGRRRRPPQGWVRQVQDRVL*

>cTP-KO4

MSTAAAAAVCPELVLRHCHPALPRHSRRQPRLPITVLLLLLLQQQQPPGPLPPPPHPQHPRLRRPRRPRPRWTPPSTASRMGSPSSRPSSMTRSRSSSSRSRPARHPSTWCSRAPTLACARR*
