## Supplemental File 5 for "Realisation of a key step in the evolution of C_4_ photosynthesis in rice by genome editing": Primer_sequences.docx

FHOx6	TGCAGCACCAGCTCTGGGCAGCGG

FHOx7	AAACCCGCTGCCCAGAGCTGGTGC

FHOx8	TGCAGAAGCACCAGCTCTGGGCAG

FHOx9	AAACCTGCCCAGAGCTGGTGCTTC

FHOx10	TGCATAATGCACGTACCGCGGCGG

FHOx11	AAACCCGCCGCGGTACGTGCATTA

FHOx90	GTGGTCTCAAATGGCGTCCAAAGGGGA

FHOx91	GTGGTCTCAAAGCTTATTTATACA

Hyg-F	CAACCAAGCTCTGATAGAGT

Hyg-R	GAAGAATCTCGTGCTTTCA
