## Supplementary figures and images for "Realisation of a key step in the evolution of C_4_ photosynthesis in rice by genome editing"

### C441062.jpg

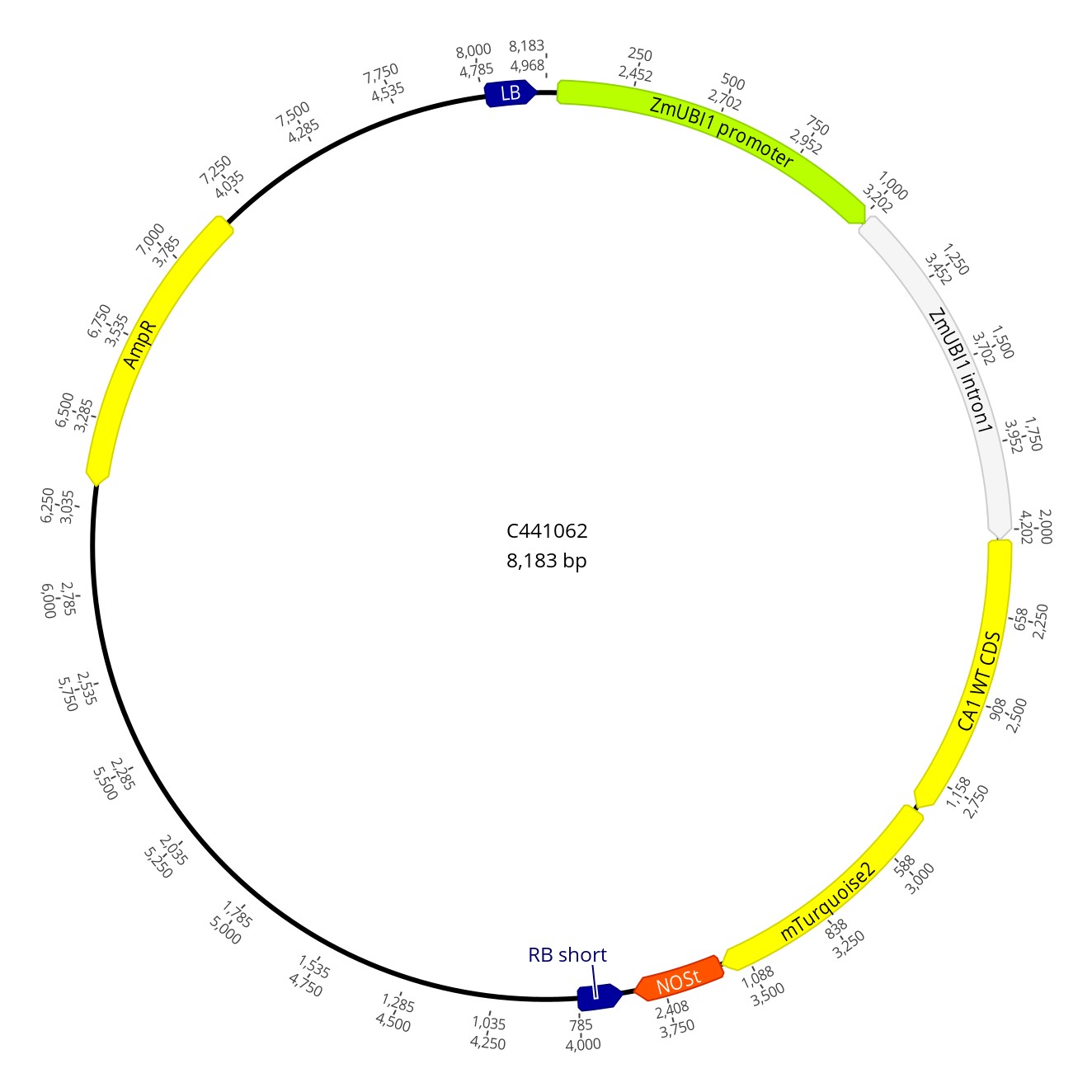

### C441064.jpg

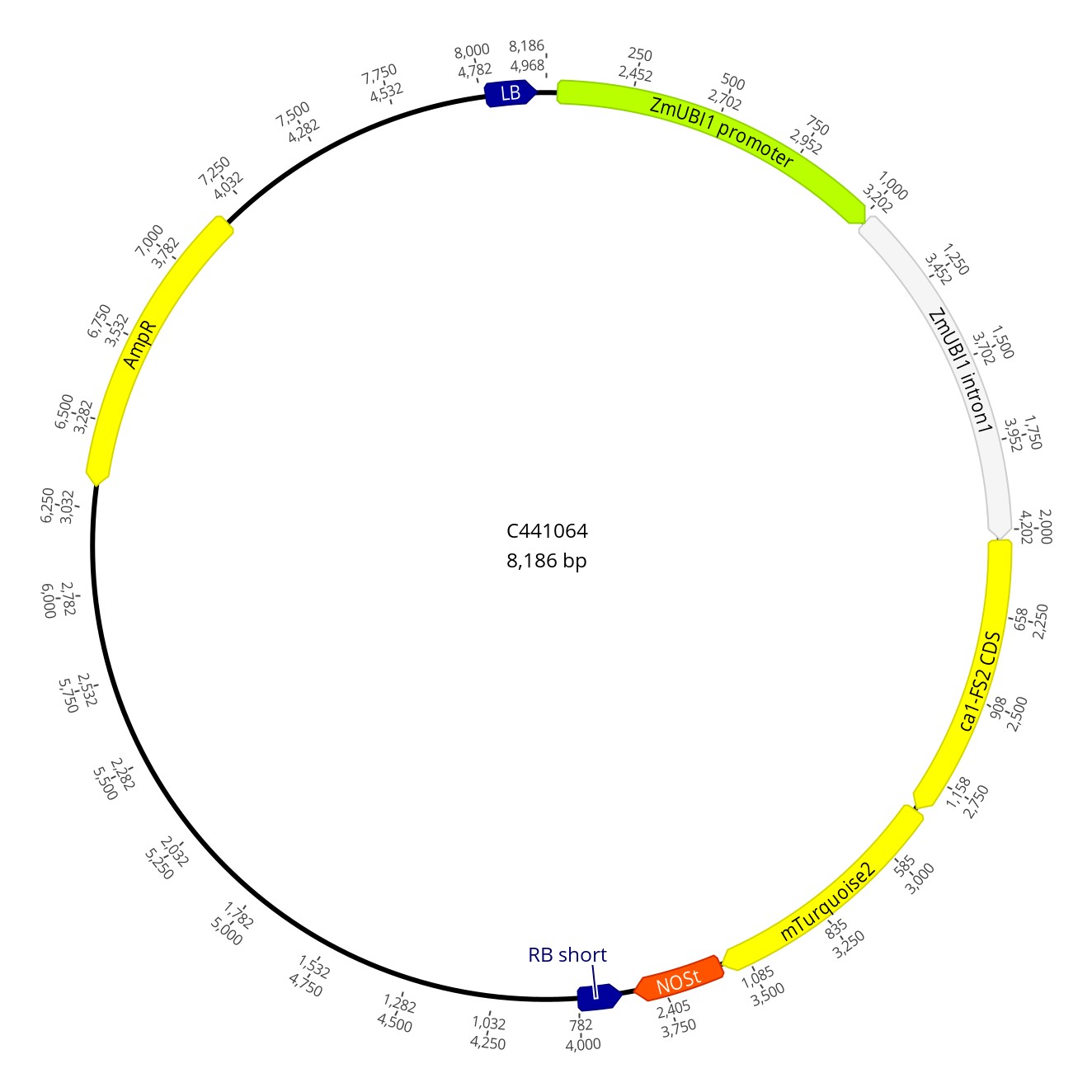

### C441065.jpg

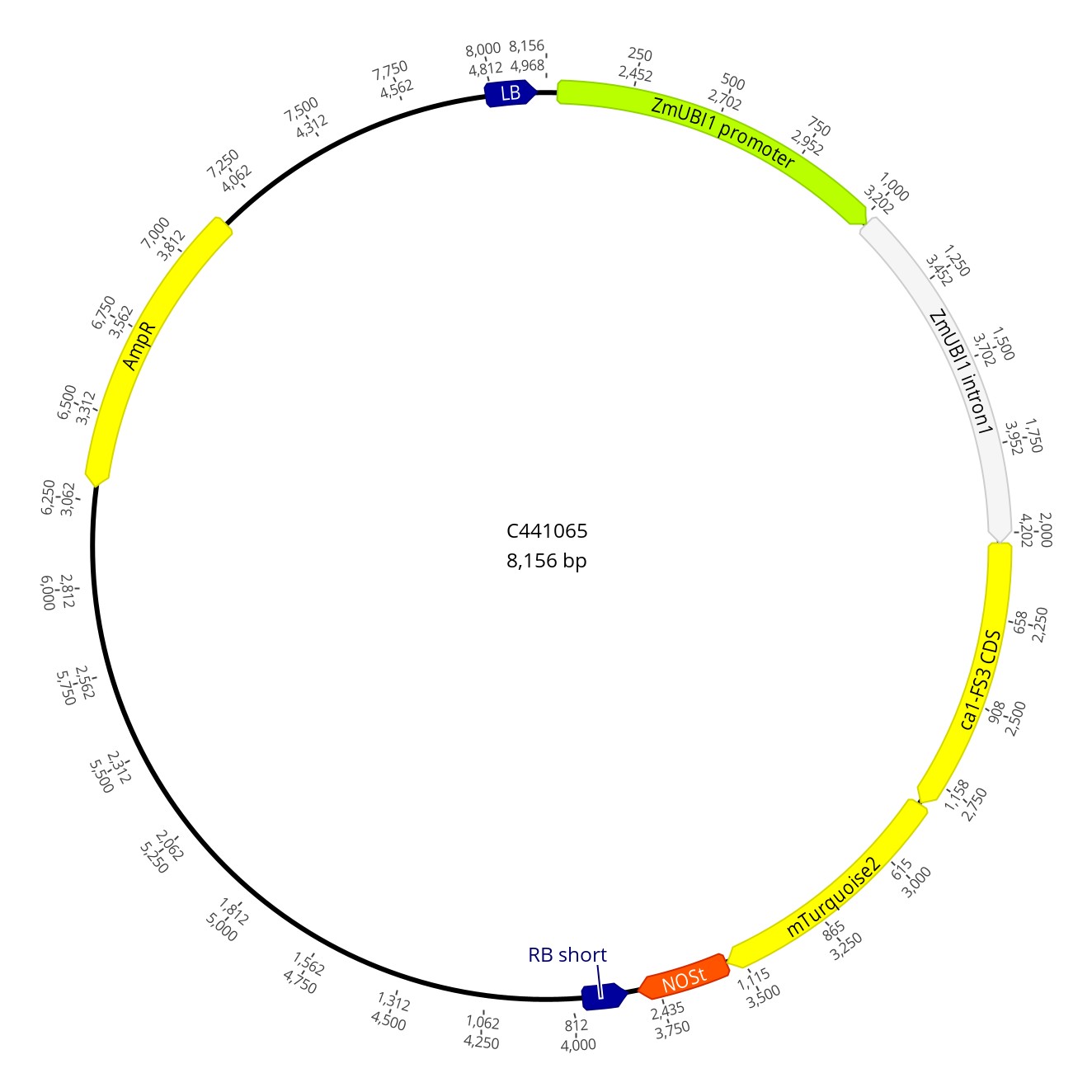

### C441067.jpg

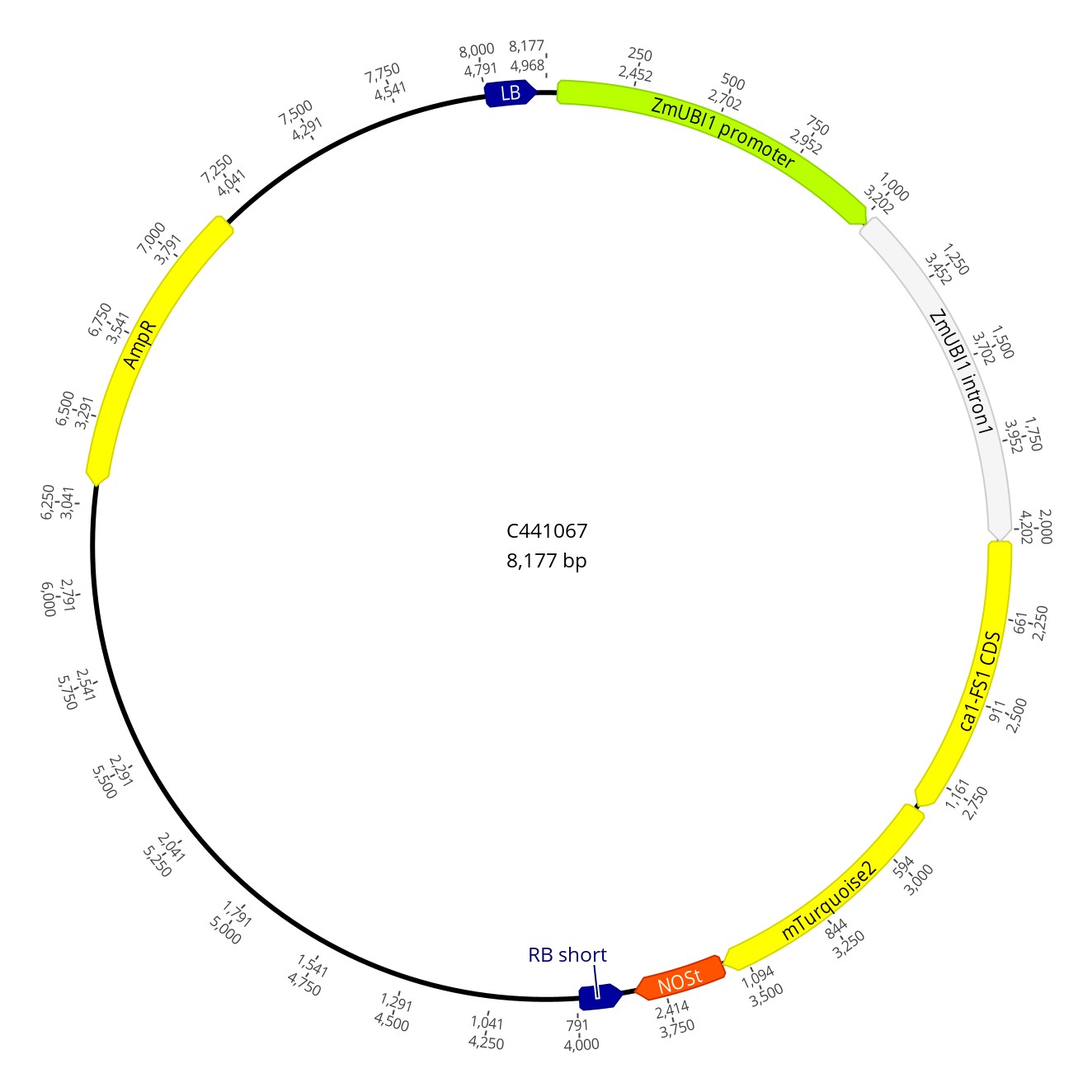

### C441068.jpg

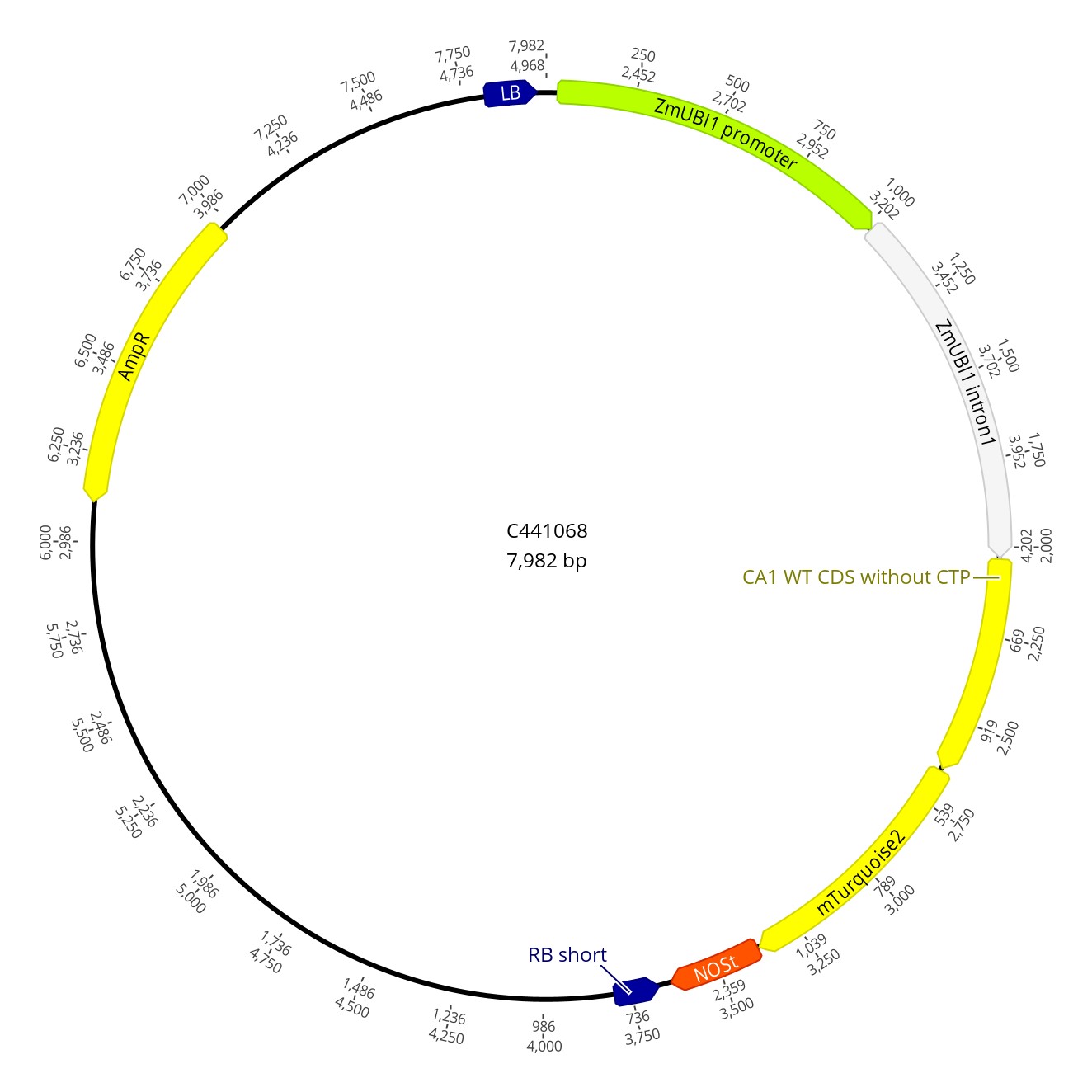

### C441069.jpg

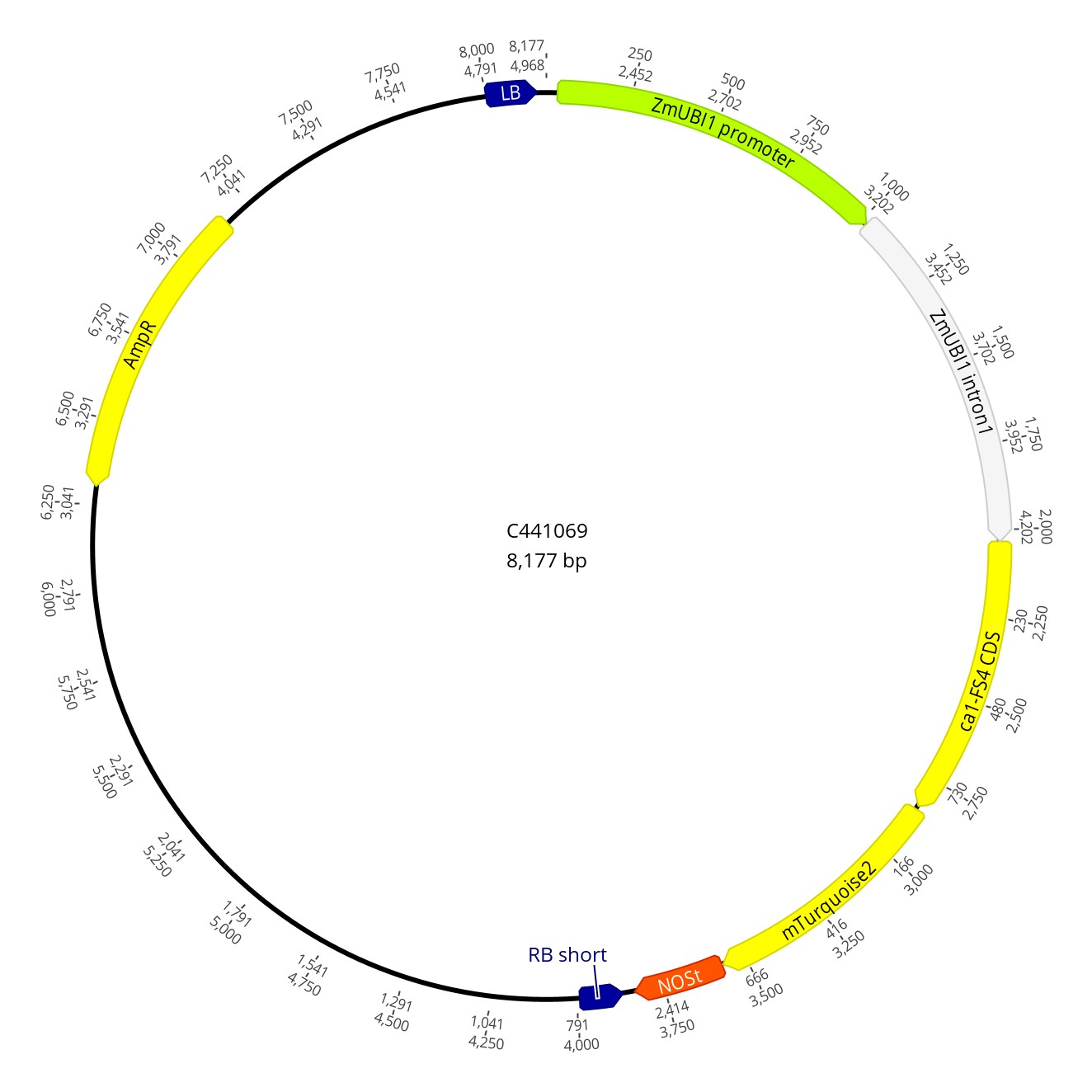

### C441070.jpg

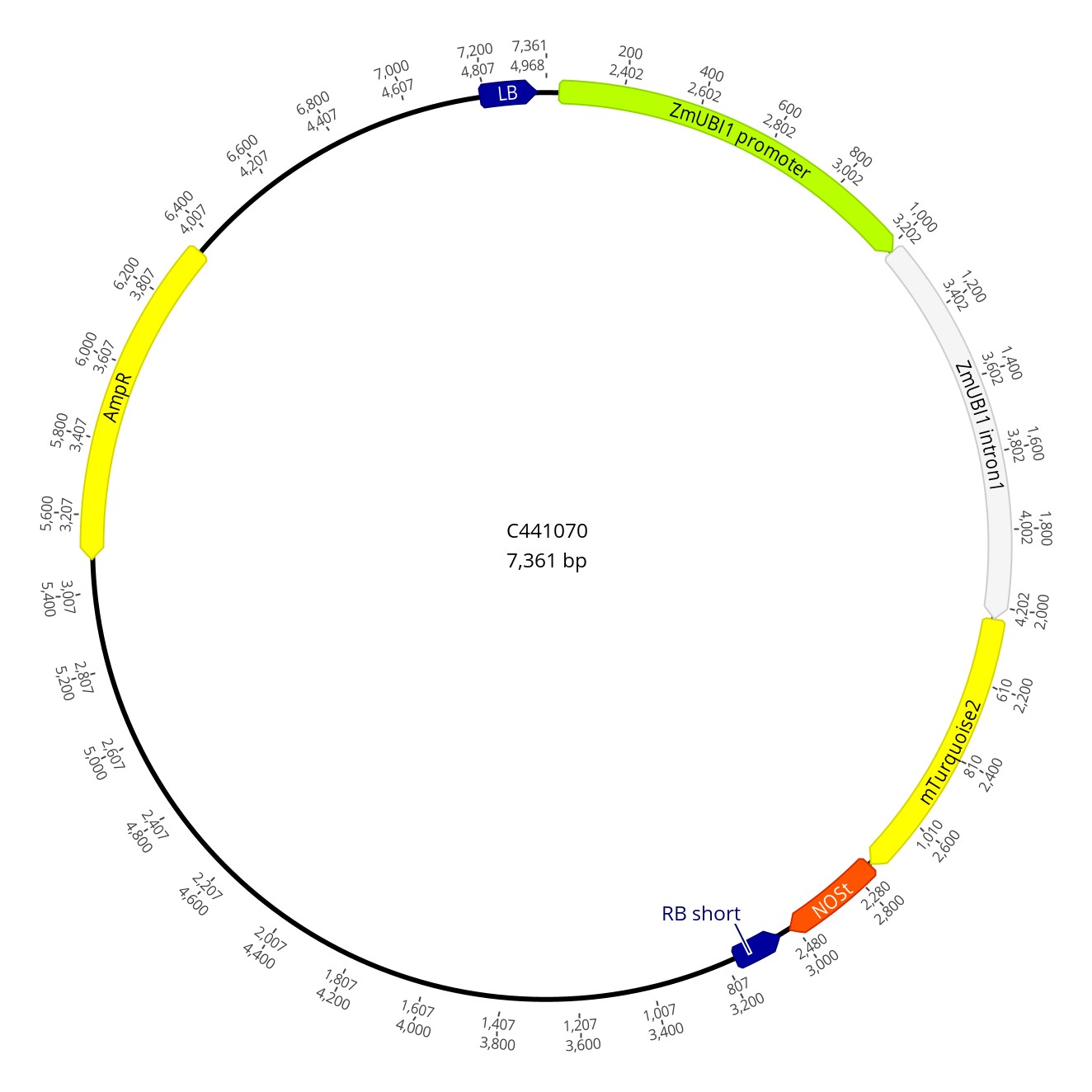

### C442001.jpg

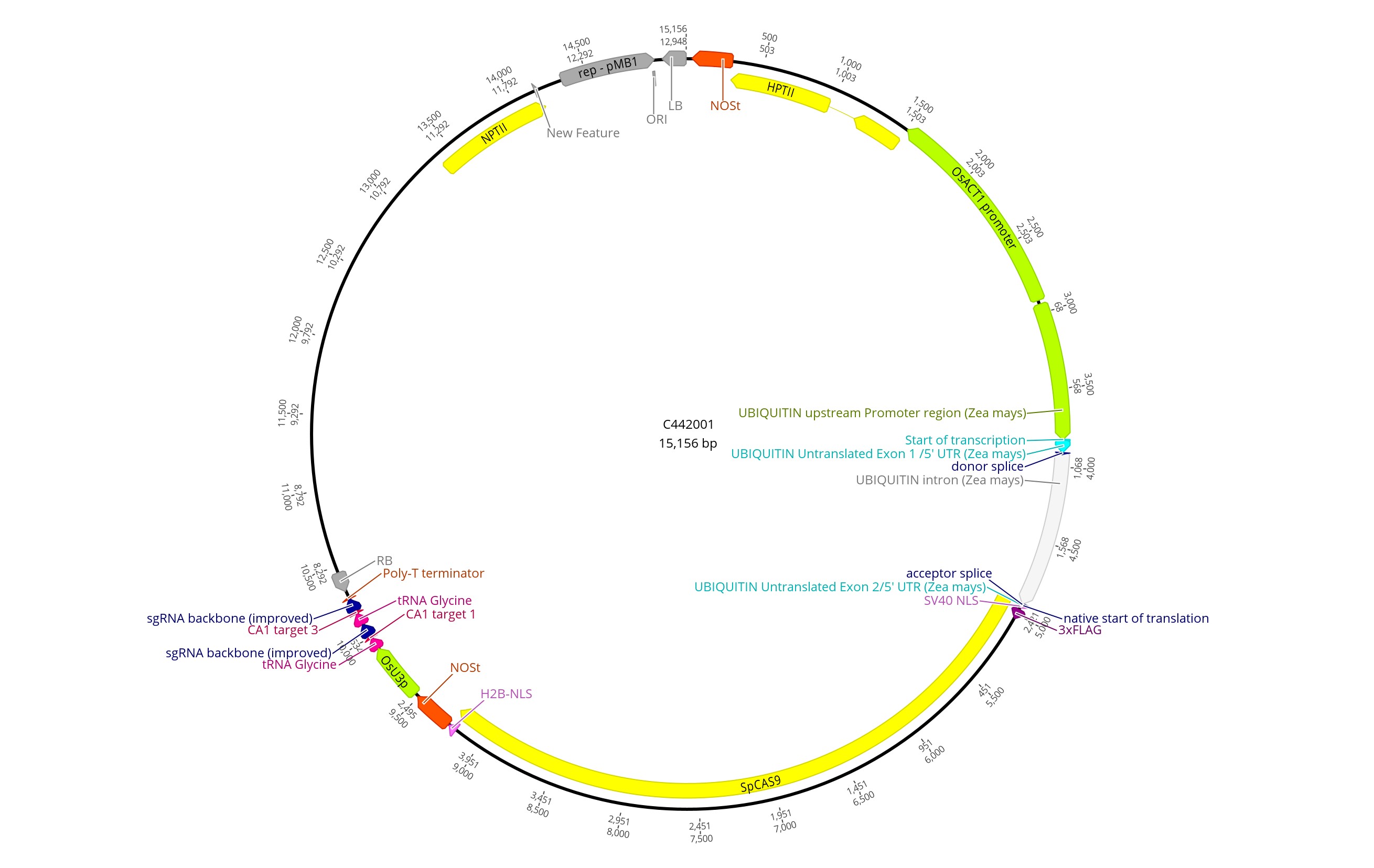

### C442002.jpg

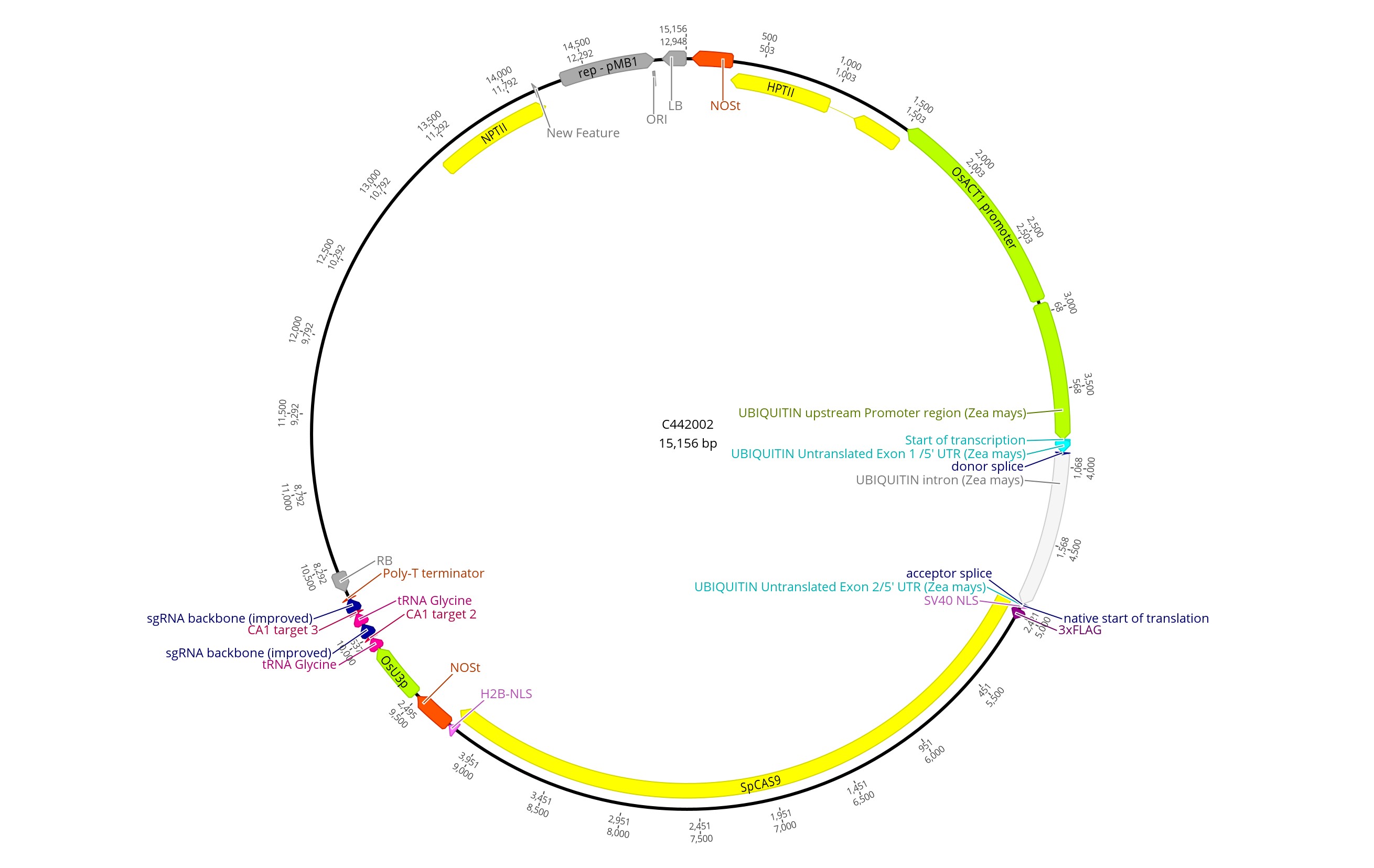
